# Identification of key residues in MERS-CoV and SARS-CoV-2 main proteases for resistance against clinically applied inhibitors nirmatrelvir and ensitrelvir

**DOI:** 10.1101/2023.12.04.569917

**Authors:** Laura Krismer, Helge Schöppe, Stefanie Rauch, David Bante, Bernhard Sprenger, Andreas Naschberger, Francesco Costacurta, Anna Fürst, Anna Sauerwein, Bernhard Rupp, Teresa Kaserer, Dorothee von Laer, Emmanuel Heilmann

## Abstract

The Middle East Respiratory Syndrome Coronavirus (MERS-CoV) is an epidemic, zoonotically emerging pathogen initially reported in Saudi Arabia in 2012. MERS-CoV has the potential to mutate or recombine with other coronaviruses, thus acquiring the ability to efficiently spread among humans and become pandemic. Its high mortality rate of up to 35 % and the absence of effective targeted therapies call for the development of antiviral drugs for this pathogen. Since the beginning of the SARS-CoV-2 pandemic, extensive research has focused on identifying protease inhibitors for the treatment of SARS-CoV-2. Our intention was therefore to assess whether these protease inhibitors are viable options for combating MERS-CoV. To that end, we used previously established protease assays to quantify inhibition of the SARS-CoV-2 and MERS-CoV main proteases. Furthermore, we selected MERS-CoV-M^pro^ mutants resistant against nirmatrelvir, the most effective inhibitor of this protease, with a safe, surrogate virus-based system, and suggest putative resistance mechanisms. Notably, nirmatrelvir demonstrated effectiveness against various viral proteases, illustrating its potential as a broad-spectrum coronavirus inhibitor. To adress the inherent resistance of MERS-CoV-M^pro^ to ensitrelvir, we applied directed mutagenesis to a key ensitrelvir-interacting residue and provided structural models.

**One-Sentence Summary:** We investigate antivirals for MERS-CoV with a pool of SARS-CoV-2 antiviral drugs and study potential resistances developing against those drugs.

## INTRODUCTION

Since the beginning of the Severe Acute Respiratory Syndrome Coronavirus 2 (SARS-CoV-2) pandemic, extensive research, governmental and industry efforts have been directed towards virus containment and therapeutic strategies (*1*, *2*). Coronaviruses generally seem very apt to first zoonotically transmit and then establish human reservoirs, as observed in human common cold coronaviruses (229E, NL63, OC43 and HKU1) (*3*). The first documented addition to human-infecting coronaviruses was SARS-CoV-1, which caused an outbreak in 2003 but was contained before becoming pandemic (*4*). Furthermore, the zoonotic and highly pathogenic Middle East Respiratory Syndrome Coronavirus (MERS-CoV) jumped into humans in 2012 and remains a source of concern, especially in Saudi Arabia and neighboring countries, as well as North America and Europe. MERS-CoV has a higher case fatality ratio (up to 35 %) than SARS-CoV-1 (11 %) and SARS-CoV-2 (*1*, *5–7*), where the global case fatalities varied depending on country from 1.7 % to 39.0 % in 2020, but stabilized at an approximate worldwide average of 0.3 % by 2022 (*8*). The substantial mortality of MERS-CoV warrants extensive research attention (*1*). Besides the potential adaptation of MERS-CoV to humans and thereby its pandemic risk, coronaviruses possess the capacity to undergo recombination (*9*). Such a recombination event between pandemic SARS-CoV-2 and endemic MERS-CoV could have unpredictable consequences. In fact, a recently described recombinant of feline and canine coronaviruses has shown a strong increase in transmissibility (*10*).

MERS-CoV sequences have been found in bats, considered the primary reservoir, and dromedary camels, acting as intermediate hosts (*11–13*). MERS-CoV is enzootically transmitted across the Arabian Peninsula and parts of Africa, occasionally spilling over to humans (**Fig. 1A**). The first documented human case emerged in Saudi Arabia in 2012, however, the exact transmission route remains unknown (*5*, *14–17*). Human-to-human transmission requires close, prolonged contact and is favored in settings lacking infection control measures. Infected camels typically suffer from mild upper respiratory tract illness, while humans are either asymptomatic or experience lower respiratory tract disease with fever, cough, dyspnoea, pneumonia. Compared to SARS-CoV-1, MERS-CoV progresses faster to acute respiratory distress syndrome and multiorgan failure. The mortality rate associated with MERS-CoV, ranging from 20 % to 40 %, may be overestimated due to under-reporting of mild or asymptomatic cases (*14*, *15*, *18*, *19*).

Ongoing MERS-CoV outbreaks in Saudi Arabia can be attributed to the absence of preventive and control measures, such as animal and human vaccines, along with the current unavailability of specific MERS-CoV treatments (*1*, *15*, *20*). However, various therapeutic options were tried in severe MERS cases, encompassing immunotherapy using convalescent plasma and intravenous immunoglobulins, antiviral agents like interferons, ribavirin, ritonavir / lopinavir, oseltamivir and alisporivir, as well as supportive therapies such as mycophenolate mofetil, chloroquine, nitazoxamide, silvestrol and glucocorticoids (*1*, *18*, *20*, *21*). Notably, most of these approaches lack studies validating their in vivo efficacy at present. Consequently, there is a compelling need for research of effective medications (*15*).

As a member of the lineage C Betacoronavirus family, MERS-CoV is a single-stranded, positive-sense RNA virus with a genome of over 30,000 nucleotides. The genome structure includes two large open reading frames (ORF1a, ORF1b) at the 5’ end, encoding the viral replication machinery. Upon MERS-CoV infection, two non-structural polyproteins (pp1a, pp1ab) are synthesized from ORF1a and ORF1b in host cells. These proteins are then cleaved by two proteases contained within them: the main protease (M^pro^) and papain-like protease (PL^pro^) (*12*, *17*, *22*). MERS-CoV-M^pro^, as the fifth non-structural protein (nsp5) of pp1a and pp1ab, processes the viral polyproteins at eleven cleavage sites, whereas PL^pro^ cleaves three sites (**Fig. 1B**). The M^pro^ structure is very conserved with chymotrypsin-like domains I and II, along with a helical domain III, leading to its third name: 3-chymotrypsin-like protease or 3CL^pro^ (*12*, *16*, *23*, *24*). To be active, M^pro^ transitions from a monomer to a homodimer (*24*). Given the critical role of these cleavage events in replication, viral proteases are promising drug targets (*2*). In fact, protease inhibitors are potent drugs to combat various viruses, including Human Immunodeficiency Virus (HIV) (*25*), Hepatitis C Virus (HCV) (*26*) and SARS-CoV-2 (*27*, *28*).

In this study, we assesed whether SARS-CoV-2-M^pro^ inhibitors also inhibit MERS-CoV-M^pro^. We evaluated the clinically approved inhibitors nirmatrelvir and ensitrelvir across several viral proteases. We found that nirmatrelvir exhibited similar efficacy against MERS-CoV-M^pro^ compared to SARS-CoV-2-M^pro^, underlining its potential use in treating patients with MERS-CoV infections. Ensitrelvir showed less activity than nirmatrelvir against MERS-CoV-M^pro^ and other tested viral proteases. Subsequently, we applied directed mutagenesis to a key ensitrelvir-interacting residue to investigate the selective resistance mechanism of MERS-CoV-M^pro^. To extend on the broad applicability of nirmatrelvir, we explored if its use in MERS-CoV-M^pro^ could provoke viral resistance development as described for SARS-CoV-2 (*29*). To that end, we used previously developed, surrogate virus-based assays (*29–31*). We thereby selected mutants against nirmatrelvir and suggest likely resistance mechanisms based on molecular modelling.

## RESULTS

### Sequence and structural relation between SARS-CoV-2 and MERS-CoV main proteases

For both SARS-CoV-2 and MERS-CoV, the structures of their respective main proteases were determined by X-ray crystallography (*24*, *32*). Initially, we compared the amino acid sequence of MERS-CoV-M^pro^ with that of SARS-CoV-2-M^pro^, revealing 51 % sequence identity (**Fig. 1C**). The tertiary structures of both main proteases (e.g. Protein Data Bank (PDB) IDs 7VH8, 5C3N) are highly similar (RMSD 253 pruned atom pairs: 0.906 Å) (**Fig. 1D**), and more conserved than the sequences. This is often the case between related proteins, as structure dictates function (*33*).

### Methods for MERS-CoV-M^pro^ activity assessment

To evaluate protease activity, we used a biosafety level 1 cellular assay based on Vesicular Stomatitis Virus (VSV) (*30*). This negative-strand RNA virus belongs to the Rhabdoviridae family and is widely studied as an oncolytic and vaccine vector, among other applications (*34*, *35*). The VSV genome encodes five essential proteins: nucleoprotein (N), phosphoprotein (P), matrix protein (M), glycoprotein (G) and polymerase (L) (*35–37*). Two chimeric VSV constructs (“On and Off”) were originally developed to regulate VSV replication with the HIV protease and then adapted to assess SARS-CoV-2-M^pro^ activity (*30*, *31*). The measurement tools rely on replication-incompetent VSV-dsRed variants missing either the P protein (VSV-ΔP-dsRed) or the L polymerase (VSV-ΔL-dsRed), which were replaced with a red fluorescent protein (dsRed). In the M^pro^-On assay (“gain-of-signal”), the intramolecular insertion of the main protease monomer into the polymerase cofactor P causes constant degradation of the artificial fusion protein P:M^pro^:P, rendering virus replication impossible. Only in the presence of protease inhibitor, the P protein remains intact, enabling VSV replication and transgene expression (**Fig. 1E**). To sensitize the construct for weaker inhibitors, one of the two cognate protease recognition sequences was knocked-out at the N-terminus by a glutamine to asparagine mutation (QtoN) (*38*). In the M^pro^-Off assay (“loss-of-signal”), the viral L polymerase was tagged N-terminally with M^pro^, enhanced green fluorescent protein (eGFP) and protease cleavage sites N-and C-terminal of M^pro^. After tag removal by proteolytic cleavage, the L protein regains its functionality, allowing VSV replication. However, adding a protease inhibitor prevents cleavage and therefore virus replication (*30*, *31*). Quantitative detection in both assays is facilitated by dsRed, whereby signal strength correlates with inhibitor activity (**Fig. 1E**).

**Figure 1.**
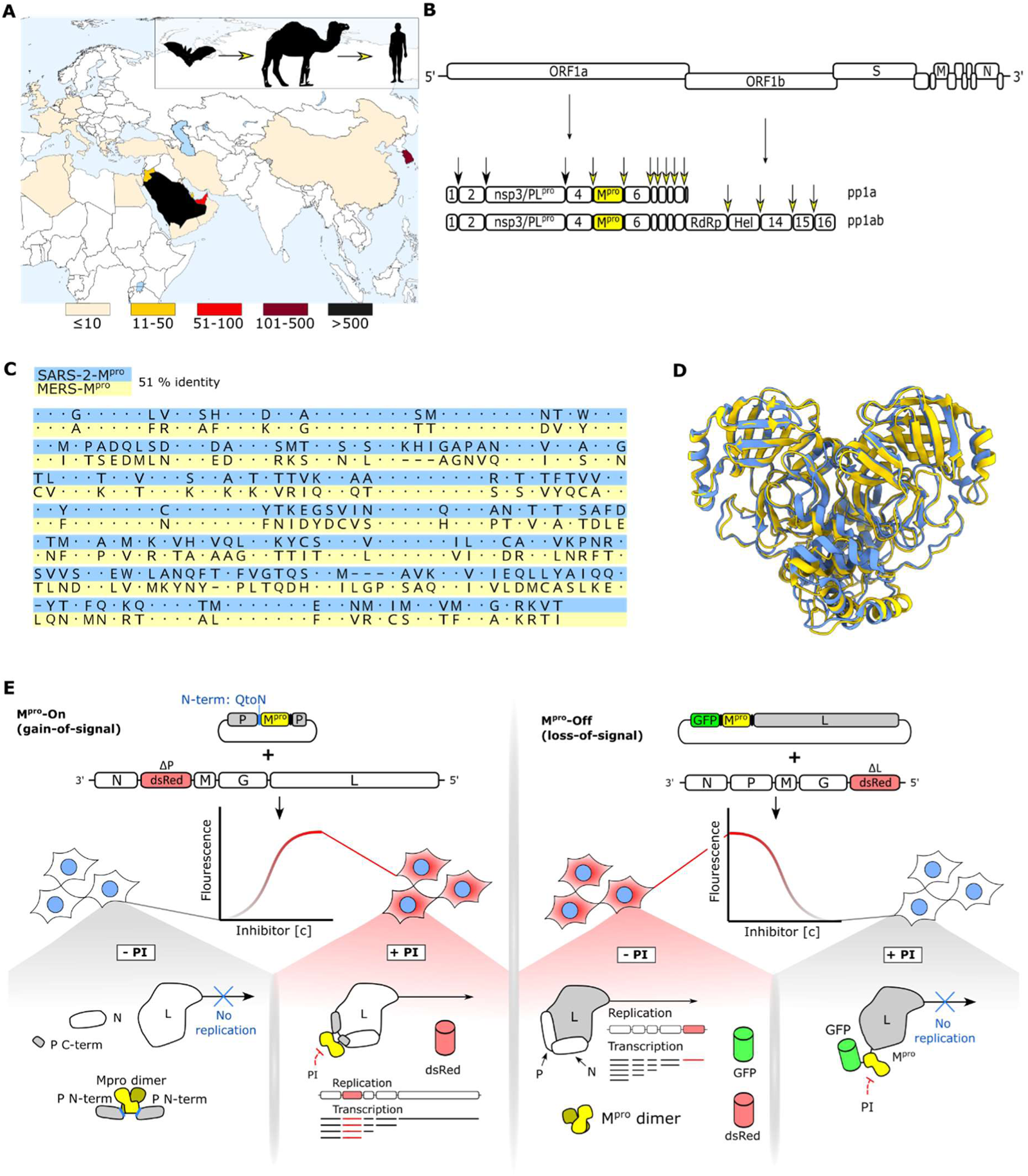
Characteristics of MERS-CoV and methods for MERS-CoV-M^pro^ activity assessment. (**A**) Epidemiology and transmission of MERS-CoV. Case counts are color coded by country from light yellow (up to 10 reported cases) to black (more than 500 reported cases) (*39*). (**B**) Schematic representation of the MERS-CoV genome. Two large polyproteins (pp1a, pp1ab) are translated from the positive-strand RNA genome. (**C**) MUSCLE sequence alignment shows 51 % identity between SARS-CoV-2 and MERS-CoV main proteases. (**D**) Superposed structures of SARS-CoV-2 and MERS-CoV main proteases show structural conservation. (**E**) Schematics of M^pro^-On and M^pro^-Off assays. In the M^pro^-On assay, VSV-ΔP-dsRed infection and P:M^pro^:P transfection result in virus replication and dsRed expression in presence of a protease inhibitor (gain-of-signal). Without inhibitor, VSV P protein auto-cleavage by M^pro^ abrogates viral replication and thereby dsRed expression. In the M^pro^-Off assay, VSV-ΔL-dsRed infection and GFP-M^pro^-L transfection without inhibitor lead to proteolytic cleavage and thereby liberated, functional L polymerase, facilitating virus replication. The addition of an inhibitor prevents cleavage and subsequently virus replication (loss-of-signal).

### Nirmatrelvir, GC376 and PF-00835231 are most effective against MERS-CoV-M^pro^

To assess antiviral activity of different protease inhibitors against MERS-CoV-M^pro^, we employed the M^pro^-On assay with both SARS-CoV-2 and MERS-CoV main proteases. We evaluated seven previously reported SARS-CoV-2-M^pro^ inhibitors (*40–45*): nirmatrelvir, ensitrelvir, GC376, PF-00835231, bofutrelvir, compound **19** and boceprevir. Nirmatrelvir, the active ingredient of Paxlovid, has received emergency use authorization for SARS-CoV-2 treatment, making it the first approved protease inhibitor for this purpose (*41*). Ensitrelvir showed potent *in vitro* antiviral activity against Omicron subvariants and is clinically authorized in Japan (*43*). GC376 was developed as a broad-spectrum 3C-protease inhibitor, and boceprevir is an approved anti-HCV drug (*40*, *46*). PF-00835231 underwent clinical trials as part of the first regimen targeting SARS-CoV-2-M^pro^ (*42*). Bofutrelvir, also known as FB2001, exhibited antiviral *in vitro* activity against several SARS-CoV-2 variants (*45*). Compound **19** was identified as a SARS-CoV-2-M^pro^ inhibitor using a structure-based virtual screening approach (*44*).

In the SARS-CoV-2-M^pro^ construct, all tested inhibitors demonstrated a dose-dependent increase in dsRed signal. Nirmatrelvir and ensitrelvir proved to be the most potent compounds, closely followed by GC376. Higher inhibitor concentrations were required for compound **19** and boceprevir to induce a signal. Notably, high inhibitor doses led to a decline in dsRed signal, particularly with bofutrelvir and compound **19**, indicating compound toxicity at these elevated concentrations (**Fig. 2A**). When assessing MERS-CoV-M^pro^, we observed fewer dsRed events and a need for higher inhibitor concentrations to achieve effective chemical inhibition than in the case of SARS-CoV-2-M^pro^. However, GC376, nirmatrelvir and PF-00835231 appeared to be potent inhibitors for MERS-CoV-M^pro^. Ensitrelvir was significantly less effective against MERS-CoV-M^pro^ than against SARS-CoV-2-M^pro^. Similar to the results of SARS-CoV-2-M^pro^, bofutrelvir and compound **19** showed toxicity at high concentrations (**Fig. 2B**).

**Figure 2.**
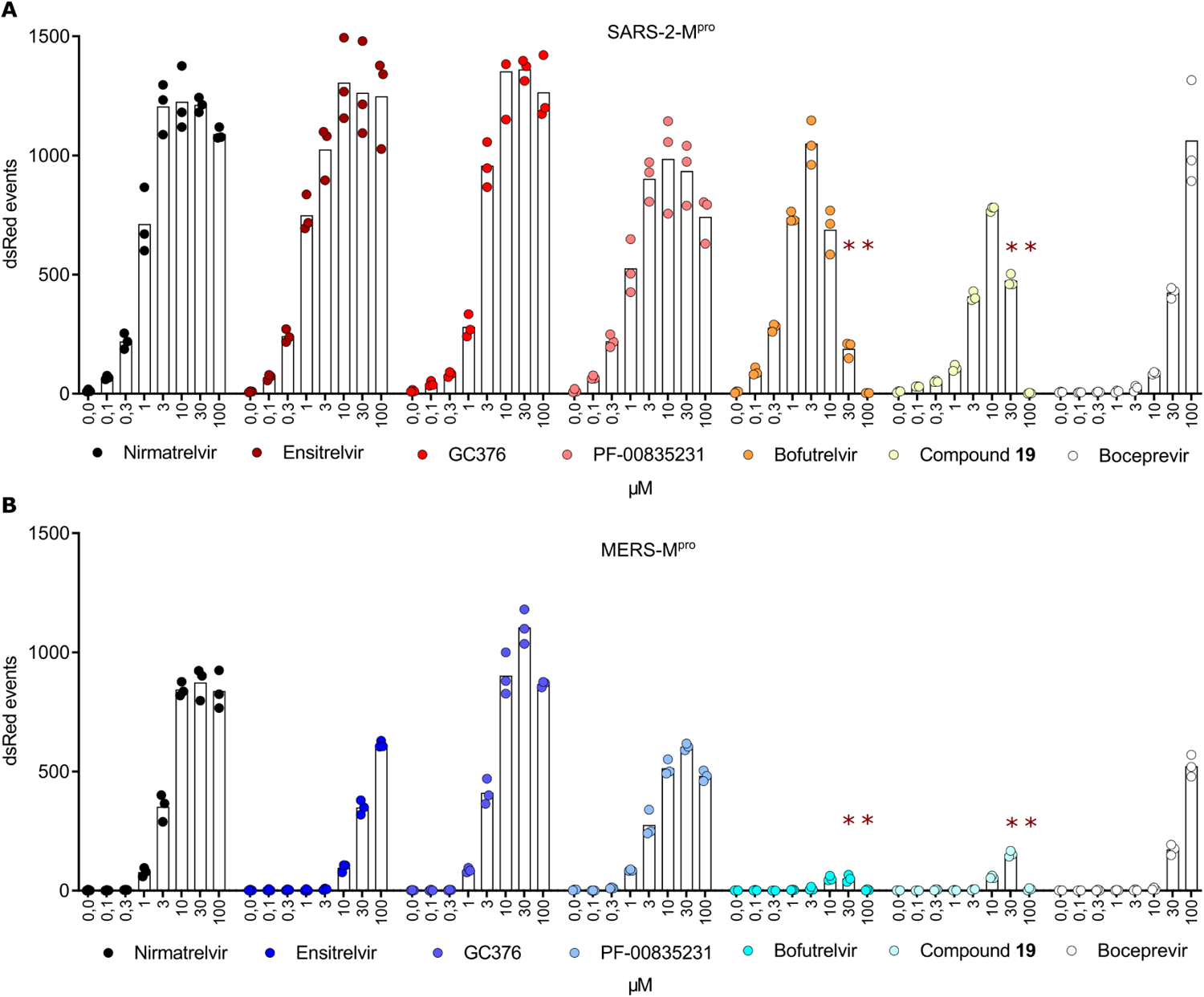
Nirmatrelvir, GC376 and PF-00835231 are most effective against MERS-CoV-M^pro^. M^pro^-On assays with protease inhibitors nirmatrelvir, ensitrelvir, GC376, PF-00835231, bofutrelvir, compound **19** and boceprevir for SARS-CoV-2-M^pro^ (**A**) and MERS-CoV-M^pro^ (**B**). Data are presented as individual data points from three biologically independent replicates per condition. Asterisks (**) indicate compound toxicity at high concentrations.

### Specific resistance mutations in SARS-CoV-2-M^pro^ and natural MERS-CoV-M^pro^ variation suggest existence of key interaction residues with the inhibitor ensitrelvir

Through mutation selection experiments in our recent work (*47*), we identified mutants that conferred selective ensitrelvir resistance in SARS-CoV-2-M^pro^. Two of them, T25A and T25N, emerged within the catalytic site, directly interacting with ensitrelvir. To confirm their selective resistance phenotypes, we performed Off assays applying nirmatrelvir and ensitrelvir. T25A exhibited increased ensitrelvir resistance and moderate resistance to nirmatrelvir compared to SARS-CoV-2-M^pro^ wild-type (wt) (**Fig. 3A**). However, T25N demonstrated resistance only against ensitrelvir (**Fig. 3B**). We previously observed that some SARS-CoV-M^pro^ mutants had delayed signal kinetics, necessitating later read-out time points for appropriate resistance assessment (*48*). To investigate whether MERS-CoV-M^pro^ also had a different kinetic than SARS-CoV-M^pro^, we first performed M^pro^-Off replication kinetics. To that end, an adapted M^pro^-Off assay was applied without adding an inhibitor. Over time, dsRed signal increased as viral replication was not disrupted. Fluorescent signals were plotted against time (hours post infection, hpi) and the TM50 metric described the time required for the curve to reach half of its maximum value when the signal plateaus (**fig. S1A**). Indeed, MERS-CoV-M^pro^ exhibited a slower kinetic, thus we opted for read-out timepoints at 70 – 80 hpi (**fig. S1B**). Subsequent M^pro^-Off assays showed moderate resistance against nirmatrelvir and strong resistance against ensitrelvir in MERS-CoV-M^pro^ wt (**Fig. 3C**). In summary, SARS-CoV-M^pro^ mutants T25A and -N as well as MERS-CoV-M^pro^ wt exhibited varying degrees of ensitrelvir-specific resistance (**Fig. 3D**).

In light of ensitrelvir’s limited effectiveness against MERS-CoV-M^pro^, we investigated potential resistance mechanisms by comparing the main proteases of SARS-CoV-2 and MERS-CoV. We first generated Maestro 2D representations of their catalytic sites, derived from X-ray structures of SARS-CoV-2-M^pro^ bound to nirmatrelvir (**Fig. 3E**), ensitrelvir (**Fig. 3F**) and MERS-CoV-M^pro^ bound to nirmatrelvir (**Fig. 3G**). We used Autodock 4 to generate an ensitrelvir-bound MERS-CoV-M^pro^ model, as this complex structure was unavailable (**Fig. 3H**). Subsequently, we compared the amino acid residues forming the catalytic site within 4 Å from nirmatrelvir and ensitrelvir, respectively. In sequence alignments, small deletions and insertions of SARS-CoV-2 and MERS-CoV main proteases led to so-called “register shifts” (**fig. S2A, B**), meaning a different numbering of structurally aligned amino acids. We annotated the stretches with identical numbering in the standard convention, i.e. residue-SARS-CoV-2-M^pro^_number_residue-MERS-CoV-M^pro^, e.g. M49L. To highlight the register shift after a relative deletion or insertion in one of the two sequences, we annotated with residue-SARS-CoV-2-M^pro^_number-SARS-CoV-2-M^pro^ / number-MERS-CoV-M^pro^_residue-MERS-CoV-M^pro^, e.g. N142 / 145C.

Within 4 Å of the nirmatrelvir binding site, a quantitative difference of three amino acid residues was observed between the two main proteases (SARS-CoV-2-M^pro^ with 21aa, MERS-CoV-M^pro^ with 18aa). Furthermore, a total of six amino acid residues were distinct between both interaction sites: M49L, N142 / 145C, H164 / 167Q, P168 / 171A, R188 / 191K and T190 / 193V (**fig. S2A, S3**). Notably, the alanine at position 171 of MERS-CoV-M^pro^ was outside the 4 Å zone (**Fig. 3E, G**). In the 4 Å zone of ensitrelvir, there was one quantitative discrepancy between SARS-CoV-2 and MERS-CoV main proteases (SARS-CoV-2-M^pro^ with 19aa, MERS-CoV-M^pro^ with 20aa). Natural amino acid variations were also present at six positions: T24S, T25M, M49L, N142 / 145C, H164 / 167Q and R188 / 191K (**fig. S2B, S4**). The serine at position 24 in MERS-CoV-M^pro^ and the methionine at position 49 in SARS-CoV-2-M^pro^ were not part of the 4 Å zone (**Fig. 3F, H**). The previously described specific ensitrelvir resistance mutants T25A and T25N (*47*) suggested the existence of key interaction residues contributing to ensitrelvir resistance. We therefore aligned the sequences of MERS-CoV and SARS-CoV-2 main proteases, focusing on the two non-conserved amino acid positions 24 and 25 in an otherwise conserved region. In SARS-CoV-2-M^pro^, positions 24 and 25 are two threonines, while in MERS-CoV-M^pro^, there are serine and methionine (**Fig. 3I**).

**Figure 3.**
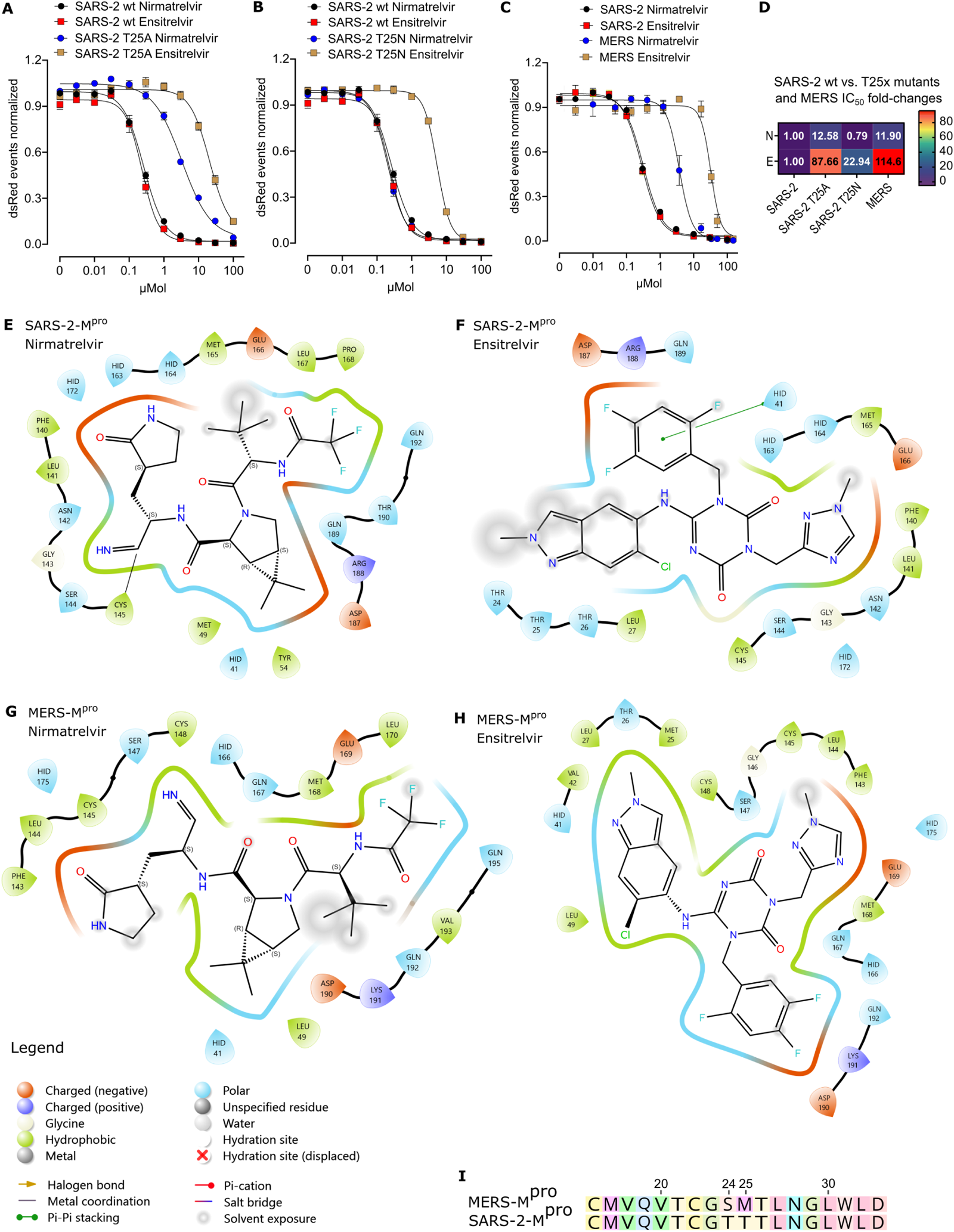
Specific resistance mutations in SARS-CoV-2-M^pro^ and natural MERS-CoV-M^pro^ variation suggest existence of key interaction residues with the inhibitor ensitrelvir. M^pro^-Off assays to assess nirmatrelvir (N) and ensitrelvir (E) efficacy in SARS-CoV-2-M^pro^ wt and SARS-CoV-2-M^pro^ ensitrelvir mutant T25A (**A**) and T25N (**B**). (**C**) M^pro^-Off assays of nirmatrelvir and ensitrelvir testing SARS-CoV-2-M^pro^ wt and MERS-CoV-M^pro^ wt. Data are presented as means of n = 4 biologically independent replicates per condition. (**D**) Heat map showing IC50 fold changes of (**A**), (**B**) and (**C**). Maestro 2D maps of SARS-CoV-2-M^pro^ catalytic site interacting with nirmatrelvir (**E**) and ensitrelvir (**F**). Interaction of MERS-CoV-M^pro^ catalytic site with nirmatrelvir (**G**) and ensitrelvir (**H**). Types of interacting residues are displayed in red (negatively charged), dark blue (positively charged), green (hydrophobic) and light blue (polar). Different binding types are also shown in the legend. Amino acid abbreviations: ARG (R), ASN (N), ASP (D), CYS (C), GLN (Q), GLU (E), GLY (G), HID (H), LEU (L), LYS (K), MET (M), PHE (F), PRO (P), SER (S), THR (T), TYR (Y), VAL (V). (**I**) Partial sequence alignment of MERS-CoV and SARS-CoV-2 main proteases, highlighting non-conserved amino acid positions 24 and 25.

### Directed mutagenesis at position 25 underlines the importance of two catalytic site loops in nirmatrelvir and ensitrelvir binding

Since MERS-CoV-M^pro^ was more resistant to ensitrelvir than to nirmatrelvir (**fig. S5A**) and T25 in SARS-CoV-M^pro^ seemed to be a hot-spot for specific ensitrelvir resistance, we investigated the potential influence of the MERS-CoV-M^pro^ methionine residue at this position by generating a “SARS-CoV-2 to MERS-CoV” mutant, T25M. However, we found no significant difference compared to SARS-CoV-2-M^pro^ wt (**Fig. 4A**). Likewise, the kinetic of T25M was similar to the wild-type (**fig. S5B**). In parallel, we attempted to resensitize MERS-CoV-M^pro^ to ensitrelvir by introducing a “MERS-CoV to SARS-CoV-2” mutation (M25T). Yet, this mutation was even more resistant to ensitrelvir (**Fig. 4B; fig. S5C**), with a mild effect on MERS-CoV-M^pro^ kinetics (**fig. S5D**). Moreover, M25T was more susceptible to nirmatrelvir than MERS-CoV-M^pro^ wt (**Fig. 4B**).

To explore the resistance of M25T, we conducted structural analyses and stability predictions. T25 in SARS-CoV-2-M^pro^ (PDB entry 7VLQ (*49*)) and M25 in MERS-CoV-M^pro^ (PDB entry 7VTC (*49*)) are located in loops made of residues 21 – 26. In both proteases, a neighboring loop is formed by residues 42 – 49 (**Fig. 4C-D**). T25 in SARS-CoV-2-M^pro^ interacts with backbone atoms of C22 and V42 in chain A (7VLQ) (**Fig. 4C**). In chain B (7VLQ), T25 interacts via hydrogen bonds with C22 and C44. In other nirmatrelvir-bound (7VH8) and apo (7ALI) structures, T25 also forms hydrogen bonds with those cysteines. In the nirmatrelvir-bound structure 8DZ2, C22, V42 and C44 hydrogen-bond with T25 in both chains. Similarly, T25 in a simulated MERS-CoV-M^pro^ M25T mutant interacted also with C44 of the neighboring loop (**Fig. 4E**). Alignment of apo (PDB ID 5C3N (*24*)) and nirmatrelvir bound (PDB ID 7VTC (*49*)) experimental MERS-CoV-M^pro^ wt structures showed a different loop arrangement. In the nirmatrelvir-bound structure, L49 is closer to H41 and interacts with nirmatrelviŕs dimethylcyclopropyl proline moiety (**Fig. 4F**). Molecular dynamic (MD) simulations were used to further investigate the impact of M25T on the flexibility of this region and particularly the movement of L49. The distance of L49 to H41 in MERS-CoV-M^pro^ M25T is similar to the nirmatrelvir-bound MERS-CoV-M^pro^ wt structure, whereas in the apo MERS-CoV-M^pro^ wt, the distance of L49 to H41 is greater (**Fig. 4G**).

To study the role of the non-conserved amino acid position 24, we also tested mutant S24T and double mutant S24T / M25T. MERS-CoV-M^pro^ wt and S24T exhibited similar susceptibilities to nirmatrelvir and ensitrelvir (**fig. S5E**). The double mutant S24T / M25T demonstrated strong resistance to ensitrelvir, whereas its sensitivity to nirmatrelvir was higher than that of wild-type (**fig. S5F, G**). Comparing SARS-CoV-2-M^pro^ wt with MERS-CoV-M^pro^ wt and its mutants directly, the added resistance to ensitrelvir by M25T and S24T / M25T was around 1,000-fold and 1,500-fold, respectively (**fig. S5H**).

**Figure 4.**
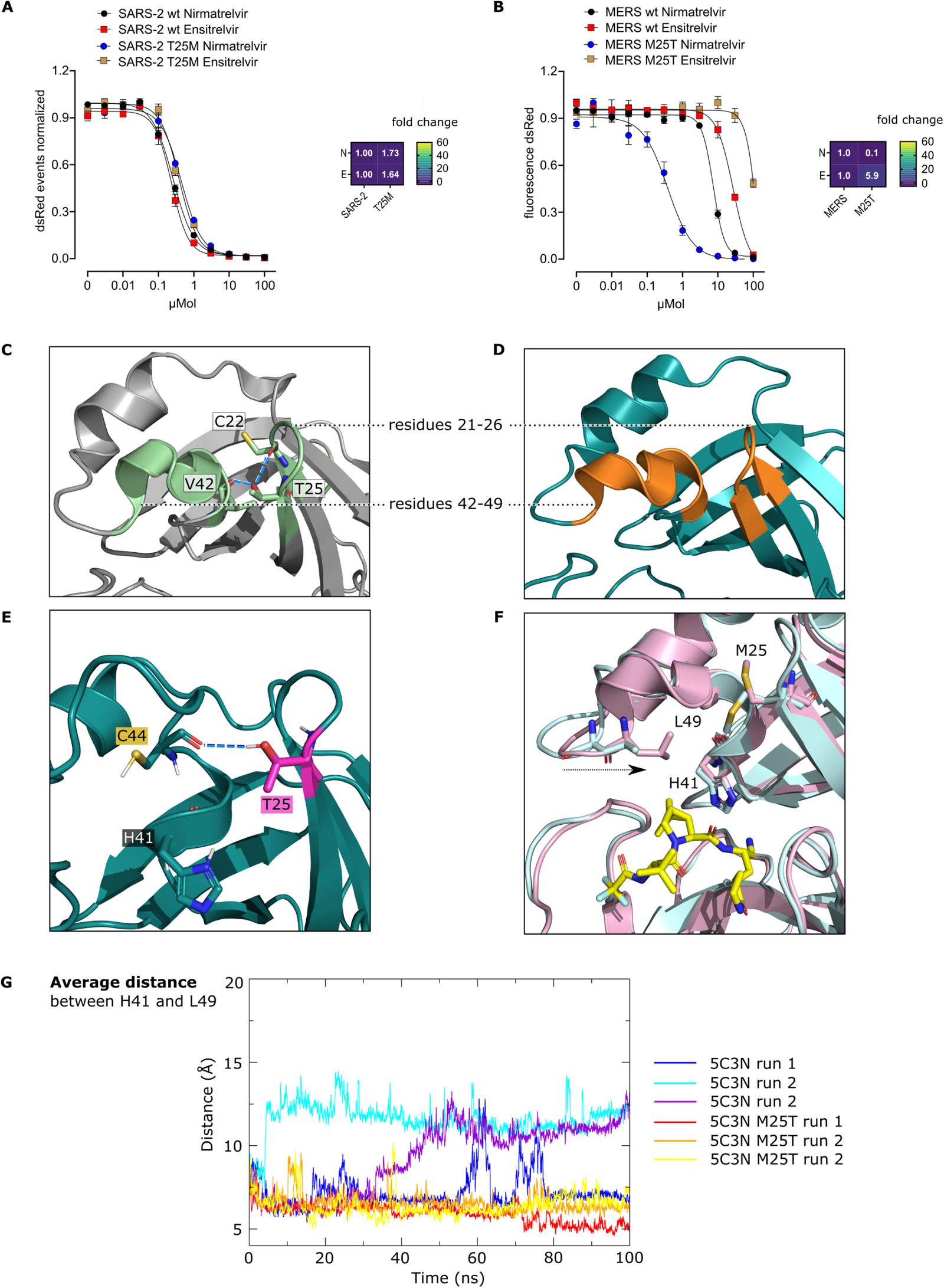
Directed mutagenesis at position 25 underlines the importance of two catalytic site loops in nirmatrelvir and ensitrelvir binding. M^pro^-Off assays and heat maps with nirmatrelvir (N) and ensitrelvir (E) testing SARS-CoV-2-M^pro^ wt and “SARS-CoV-2 to MERS-CoV” mutant T25M (**A**) and MERS-CoV-M^pro^ wt and “MERS-CoV to SARS-CoV-2” mutant M25T (**B**). (**C**) Two neighboring loops formed by residues 21-26 and 42-49 in SARS-CoV-2-M^pro^ (PDB entry 7VLQ (*49*)). T25 in SARS-CoV-2-M^pro^ chain A forms hydrogen bonds (dashed blue lines) with C22 and V42 backbones. (**D**) Analogous loops in MERS-CoV-M^pro^ (PDB entry 7VTC (*49*)). (**E**) In the structural analysis of the M25T mutant, the threonine (pink sticks) forms an additional hydrogen bond (dashed blue line) with the C44 backbone (petrol). (**F**) Comparison of apo (light cyan, PDB entry 5C3N (*24*)) and nirmatrelvir (yellow sticks) bound experimental MERS-CoV-M^pro^ wt (light pink, PDB entry 7VTC (*49*)) structures highlight the conformational changes of residues 21 – 26 and 42 – 49 upon ligand binding. As indicated with the black arrow, L49 moves closer to H41, and M25 adopts a different side chain conformation. (**G**) The average distance between H41 and L49 observed in three MD simulation runs of MERS-CoV-M^pro^ wt (blue, turquoise and purple) and M25T (red, orange and yellow) shows a closer proximity of these two residues in the mutant, similar as in MERS-CoV-M^pro^ wt bound to nirmatrelvir **(E).**

### Covalent nirmatrelvir inhibits a broader spectrum of viral proteases than non-covalent ensitrelvir

Considering the different nirmatrelvir and ensitrelvir efficacy against MERS-CoV-M^pro^, we performed On assays to evaluate inhibitor potency with proteases of several coronaviruses including SARS-CoV-2, SARS-CoV-1, MERS-CoV, HKU9, NL63, 229E, MHV, and poliovirus (**sequences S1-7**). We found that nirmatrelvir had a strong effect against SARS-CoV-2 and SARS-CoV-1, a moderate effect against MERS-CoV and HKU9, a mild effect against NL63 and 229E and no effect against MHV and polio proteases (**Fig. 5A**). Ensitrelvir was as effective against SARS-CoV-2 and SARS-CoV-1 as nirmatrelvir, less effective against MERS-CoV and HKU9 and not effective against NL63, 229E, MHV and polio proteases (**Fig. 5B**). To assess the likely correlation of inhibitor potency to amino acid sequence variation, we compared each of the proteases to that of SARS-CoV-2 with MUltiple Sequence Comparison by Log-Expectation (MUSCLE). SARS-CoV-2 and SARS-CoV-1 main proteases share high sequence identity of 96 %, while SARS-CoV-2 and MERS-CoV main proteases share 51 % identity (**Fig. 5C, alignment S1, 2**). However, other well-studied parts of the viral genome are not as conserved, for example the spike protein, having only 76 % sequence identity between SARS-CoV-2 and SARS-CoV-1 and 29 % between SARS-CoV-2 and MERS-CoV (**alignment S3, 4**).

Furthermore, we superimposed the X-ray structures available in the PDB for the tested viral proteases and found strong structural conservation within each viral protease bound to different inhibitors and their apo (unbound) structure, as well as conservation between the different proteases (**Fig. 5D, table S1**). To explore finer structural differences, which were not apparent by the backbone alignment, we docked nirmatrelvir and ensitrelvir into proteases of SARS-CoV-2, SARS-CoV-1, MERS-CoV, HKU9, 229E, MHV and poliovirus. HKU9-M^pro^ was modelled by I-TASSER as the solved structure was not available (*50*). We compared the docking models that achieved the highest docking score with nirmatrelvir or ensitrelvir to solved SARS-CoV-2-M^pro^ structures bound to the respective inhibitor (**Fig. 5E**). We then calculated the inhibition constants from the estimated free energies in nM for both compounds against the structures and HKU9-M^pro^ model, considering nirmatrelvir’s covalent binding (**Fig. 5E**). These virtual inhibition constants largely reflect the experimental data, apart from 229E-M^pro^, which displayed a low inhibition constant comparable to that of MERS-CoV-M^pro^.

**Figure 5.**
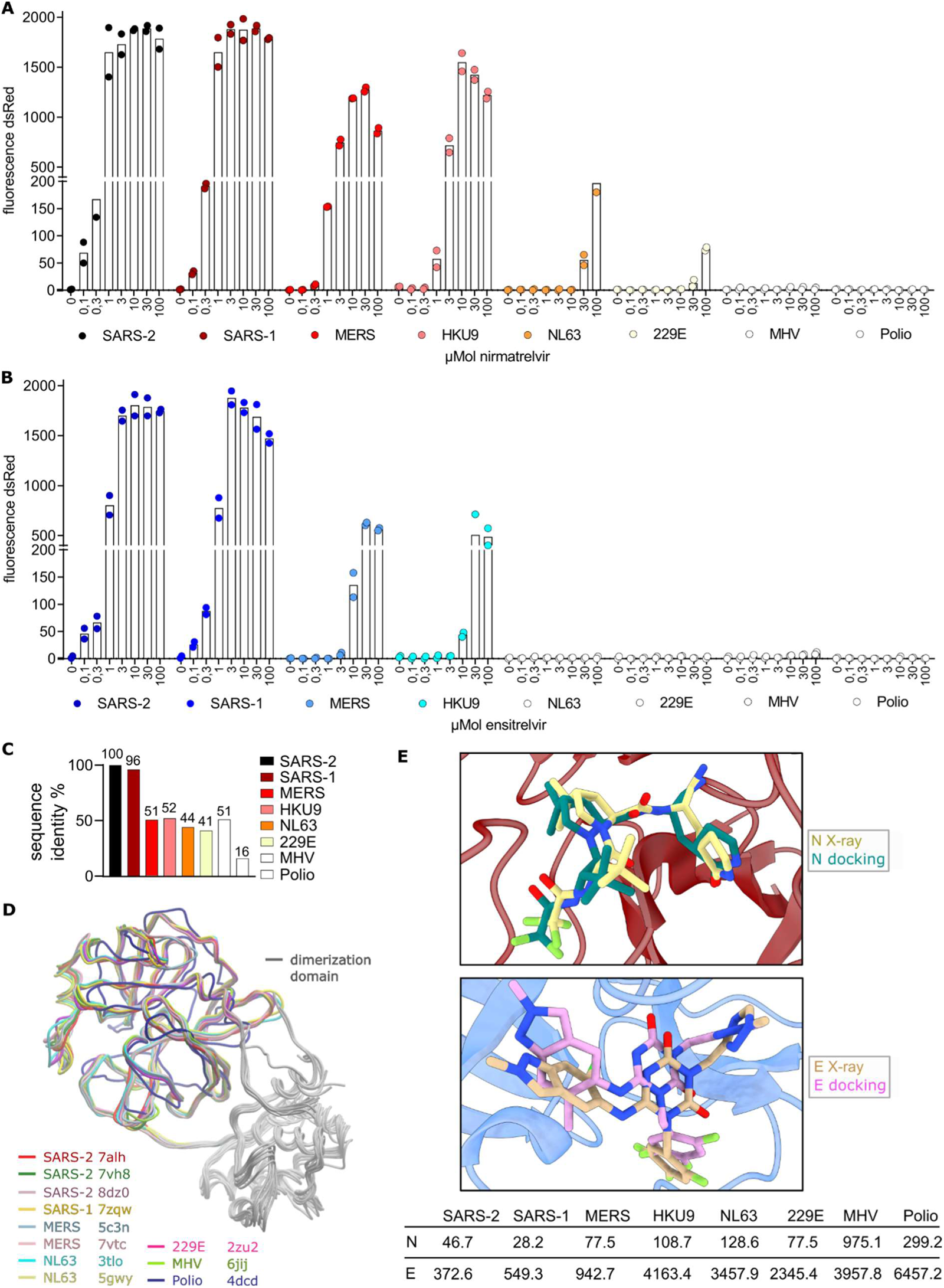
Covalent nirmatrelvir inhibits a broader spectrum of viral proteases than non-covalent ensitrelvir. M^pro^-On assays for proteases of SARS-CoV-2, SARS-CoV-1, MERS-CoV, HKU9, NL63, 229E, MHV and poliovirus with protease inhibitors nirmatrelvir (**A**) and ensitrelvir (**B**). (**C**) Comparison of sequence identities of the different viral proteases to SARS-CoV-2-M^pro^. (**D**) Structural overlay of SARS-CoV-2, SARS-CoV-1, MERS-CoV, NL63, 229E, MHV and poliovirus proteases. (**E**) Docking results of nirmatrelvir (N) and ensitrelvir (E) compared to solved X-ray structures with nirmatrelvir (PDB ID: 8DZ2) and ensitrelvir (PDB ID: 8DZ1) as well as calculated inhibition constants against a panel of viral proteases.

### Selecting and mapping nirmatrelvir resistant MERS-CoV-M^pro^ mutants

Nirmatrelvir demonstrated effectiveness against a variety of viral proteases, such as MERS-CoV-M^pro^. Given its potential use in treating MERS-CoV infections, we wanted to predict mutations of MERS-CoV-M^pro^ when exposed to nirmatrelvir. To that end, we generated two VSV variants (VSV-MERS-M^pro^) with seven and eight amino acid cleavage site lengths (7aa, 8aa) based on our previously developed mutation selection tool for SARS-CoV-M^pro^ (*29*) (**Fig. 6A**). Both variants showed similar susceptibilities to nirmatrelvir (**Fig. 6B**). We chose the 7aa variant for subsequent selection experiments with nirmatrelvir and exposed it to suboptimal nirmatrelvir concentrations, leading to continuous selection pressure through M^pro^ inhibition. After five passages of increasing nirmatrelvir doses, we selected mutants capable of replicating more effectively than the wild-type in the presence of nirmatrelvir (**Fig. 6C**).

Upon sequencing, all obtained mutants were mapped to the reference MERS-CoV-M^pro^ sequence using Geneious Prime (**Fig. 6D**). Overall, we identified 74 distinct, non-synonymous point mutations that were either alone in one sample or in combination (**table S2**). These mutations were categorized based on their location as catalytic site (S142G, S142R, S147Y, A171S), near-catalytic site (S24I, Y121F / N, K140E, G186S / C / V, F188L, D200G / Y), cleavage sites (L3S, M6K, I300K) or as allosteric mutants. We visualized mutations within the 3D dimer structure of MERS-CoV-M^pro^ (**Fig. 6E**). Some residue positions were more affected by mutations than others as shown by coloring the original residues from light blue to black. A heat map represented recurring mutations across different samples, with colors light blue (one occurrence) to dark blue (up to nine occurrences) (**Fig. 6F**).

**Figure 6.**
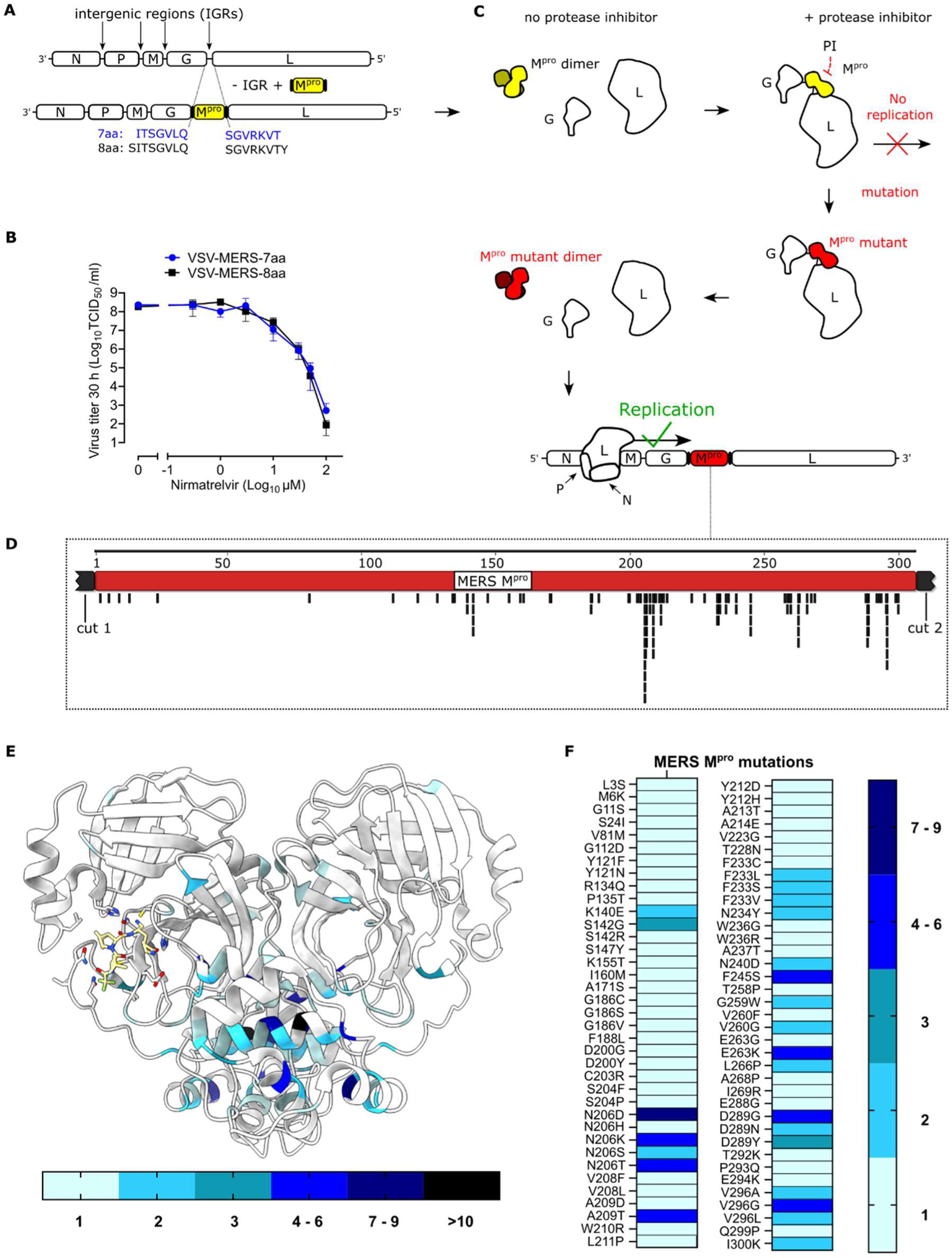
Selecting and mapping nirmatrelvir resistant MERS-CoV-M^pro^ mutants. **(A)** Schematic of generating two VSV-MERS-M^pro^ viruses (7aa, 8aa) by replacing the intergenic region (IGR) between G and L with MERS-CoV-M^pro^. G: VSV glycoprotein, L: VSV polymerase. (**B**) Nirmatrelvir dose responses of VSV-MERS-M^pro^ (7aa, 8aa). (**C**) Mutation selection tool based on resistance passages involving increasing nirmatrelvir concentrations in each passage. M^pro^ wt is highlighted in yellow, mutant M^pro^ in red. (**D**) Selected mutants mapped to reference MERS-CoV-M^pro^ sequence using Geneious Prime. (**E**) Visualization of specific mutants within the 3D dimer structure of MERS-CoV-M^pro^ using ChimeraX. The frequency of mutations at certain residues is shown by a scale from light blue (one mutation) to black (> ten mutations). (**F**) Heat map showing the number of specific MERS-CoV-M^pro^ mutations from different samples of VSV-MERS-M^pro^ nirmatrelvir selections. The frequency of specific mutations is shown by a scale from light blue (one mutation) to dark blue (seven to nine mutations).

### Characterization of nirmatrelvir selected MERS-CoV-M^pro^ mutants

We chose four single mutants within the catalytic site (S142G, S142R, S147Y, A171S) and confirmed their nirmatrelvir resistance by reintroducing them into the MERS-CoV-M^pro^-Off assay. Optimal signal measurement timepoints were determined through M^pro^-Off kinetics (**fig. S6A**). Among the mutants, S147Y showed the best kinetic profile, while A171S displayed the second fastest kinetic. We therefore tested MERS-CoV-M^pro^ wt (**Fig. 7A**) and these two mutants with nirmatrelvir, ensitrelvir, PF-00835231 and GC376. S147Y exhibited strong resistance to nirmatrelvir, milder resistance to PF-00835231 and GC376, and also increased inherent ensitrelvir resistance of MERS-CoV-M^pro^ (**Fig. 7B**). A171S had only a mild impact on all four inhibitors (**Fig. 7C**). Furthermore, we quantified nirmatrelvir and ensitrelvir resistance of S142G and S142R at a late timepoint (96 hpi) due to their slow kinetics. Both mutants showed increased resistance to nirmatrelvir, with S142G having a mild (**fig. S6B, D**) and S142R a strong effect (**fig. S6C, D**). They also had reduced activity against ensitrelvir. However, due to the late timepoint, resistance phenotypes might have been confounded by inhibitor degradation and signal plateauing.

The favorable kinetic, strongest resistance to inhibitors among the tested mutants (**Fig. 7D**), and the proximity of S147Y to the catalytically active cystein C148 prompted us to investigate the structural impact of this mutation through molecular modelling of the M^pro^ dimer (**Fig. 7E**). Tyrosine contains a benzyl ring, which requires more space than the hydroxyl group of serine. Therefore, tyrosine clashes with the lactam ring that is part of nirmatrelvir (**Fig. 7F**). To gain further insight into nirmatrelvir’s positioning, covalent docking analysis using Glide (*51–54*) was conducted with both MERS-CoV-M^pro^ wt and mutant structures. While the experimentally confirmed binding pose of nirmatrelvir could be reproduced with the wild-type structure, the S147Y mutation blocked the binding mode of nirmatrelvir, pushing its lactam ring upwards (**Fig. 7F**). S142 lies in the dimerization interface of the two MERS-CoV-M^pro^ protomers and forms a hydrogen bond with Q299 of the other protomer, thus playing a crucial role in dimer formation (**Fig. 7G**). The in-silico mutation to glycine (S142G) disrupted this hydrogen bond and increased loop flexibility. Similar to a previous study (*48*), the impact of mutations on the stability of specific protein conformations as well as the affinity of the functional dimer was assessed using computational protein design tools BioLuminate (*55–58*) and Osprey (*59–62*). BioLuminate generated a model of the mutated structure and calculated a Δ stability value, indicating changes in mutant stability compared to the wild-type. Negative values suggested increased and positive values decreased stability, respectively. Osprey calculated so-called Log_10_ K* scores for both the input and the mutated structure, with a decrease in score indicating a loss of affinity. Consistent with structural analysis and slower kinetics, S142G showed a Δ stability value of +10.88 kcal/mol, indicating reduced stability, and decreased affinity of the dimer (Log_10_ K* wt = 5.93 vs. Log_10_ K* S142G = 2.19). Structural analysis of S142R also revealed a loss of the hydrogen bond, and the larger size of arginine compared to serine may lead to steric clashes between the protomers (**Fig. 7G**). S142R showed even lower stability (Δ stability value of +38.08 kcal/mol) as S142G and Osprey calculations also predicted a reduced affinity, with a Log_10_ K* score of -443,48 compared to the wild-type (Log_10_ K* = 5.93). A171S showed increased affinity to nirmatrelvir and a mild destabilization of the apo structure (**fig. S6E**).

Lastly, we used a biochemical assay to confirm the resistance of MERS-CoV-M^pro^ S147Y based on the fluorogenic M^pro^ substrate Ac-Abu-Tle-Leu-Gln-MCA. M^pro^ cleavage of this substrate leads to a continuous increase in fluorescence, while adding an inhibitor prevents that increase (**fig. S6F**). MERS-CoV-M^pro^ S147Y was more resistant to the inhibitor nirmatrelvir than the purified MERS-CoV-M^pro^ wild-type protein, leading to a shift of the signal increase at higher inhibitor concentrations (**fig. S6G**).

**Figure 7.**
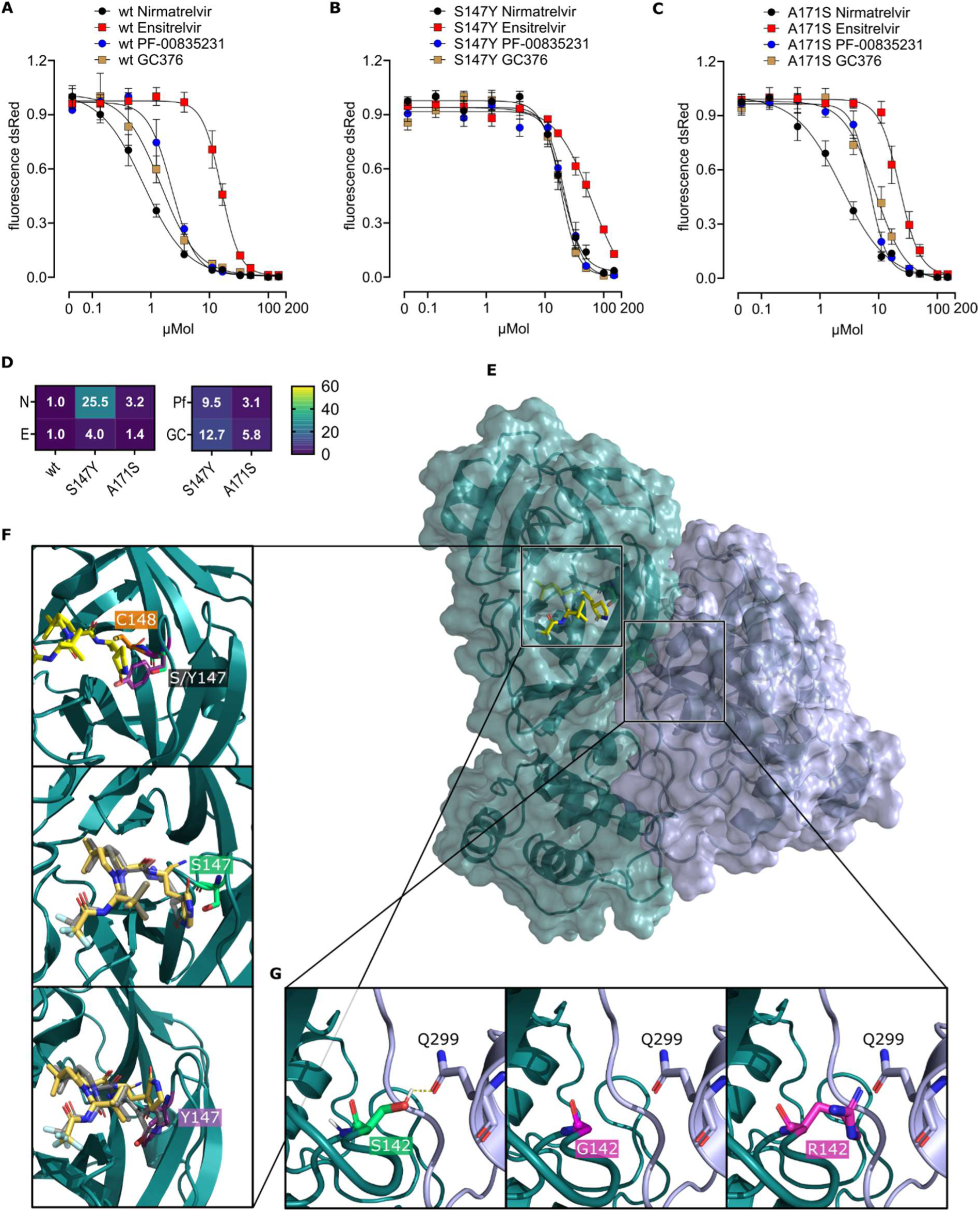
Characterization of nirmatrelvir selected MERS-CoV-M^pro^ mutants. M^pro^-Off assays to assess the susceptibility to protease inhibitors nirmatrelvir (N), ensitrelvir (E), PF-00835231 (Pf) and GC376 (GC) in MERS-CoV-M^pro^ wt (**A**) and mutants S147Y (**B**) and A171S (**C**). Data are presented as means of n = 4 biologically independent replicates per condition. (**D**) Heat map showing IC50 fold changes of (**A**), (**B**) and (**C**). (**E**) Overview of the MERS-CoV-M^pro^ homodimer bound to nirmatrelvir (yellow sticks), with two protomers colored petrol and light violet, respectively (PDB entry 7VTC (*49*)). (**F**) Structural analysis of the S147Y mutation. Upon binding to MERS-CoV-M^pro^, nirmatrelvir is positioned in close proximity to S147 (lime green). In the S147Y mutation, the tyrosine side chain (purple sticks) occupies the binding site of the nirmatrelvir lactam ring causing a clash with the inhibitor. The catalytic C148 is depicted as orange sticks. Re-docking nirmatrelvir into MERS-CoV-M^pro^ wt revealed a similar binding pose between docked (gold) and experimentally determined (grey) nirmatrelvir. Upon mutation, the nirmatrelvir lactam ring shifts upwards to prevent clashes with the Y147 side chain (purple sticks). (**G**) Structural analysis of S142G and S142R mutations. Residue S142 (lime green) on protomer 1 (petrol) forms a hydrogen bond (yellow dashed line) with Q299 on protomer 2 (light violet). The S142G or S142R mutations (pink) result in a loss of this hydrogen bond. Moreover, the larger arginine side chain causes a steric clash with Q299, thus requiring conformational changes to the interaction interface.

## DISCUSSION

In this study, we explored the effectiveness of SARS-CoV-2-M^pro^ inhibitors for MERS-CoV-M^pro^ and described the broad efficacy of nirmatrelvir. Furthermore, we identified key residues in SARS-CoV-2 and MERS-CoV main proteases associated with selected resistance to nirmatrelvir and a natural low response to ensitrelvir, providing their putative resistance mechanisms.

Human coronaviruses (CoV), first characterized in the 1960s, possess the largest genome among all known RNA viruses (*63*). The error-prone nature of RNA polymerases and the generation of subgenomic RNAs facilitate adaptability through mutations and homologous recombination, thereby increasing the likelihood of cross-species transmission and the emergence of novel CoV strains (*63–65*). Although coronavirus polymerases have some proof-reading activity (*66*), they exhibit rapid evolutionary dynamics, as exemplified by the SARS-CoV-2 pandemic. Especially the spike protein evolved quickly and defined new variants, in contrast to other parts of the viral genome, such as the main protease, which seemed to be more conserved and the penalty of mutations higher (*67*).

We therefore assessed the potential cross-viral-species applicability of licensed protease inhibitors, nirmatrelvir and ensitrelvir, against seven relevant coronaviral main proteases (SARS-CoV-2, SARS-CoV-1, MERS-CoV, HKU9, NL63, 229E, MHV) and the poliovirus protease. SARS-CoV-1 (Betacoronavirus lineage B) and MERS-CoV (Betacoronavirus lineage C) are known to be enzoonotic in bats and other species such as civets or camels with occasional spillovers into human populations (*63*, *68*). Similarly, SARS-CoV-2 (Betacoronavirus lineage B) has most likely been transmitted zoonotically from bats (*69–74*). Given that bats are a frequent source of dangerous viruses, we also tested the bat coronavirus HKU9-M^pro^ (Betacoronavirus lineage D) (*6*, *75*). Human CoV NL63 (Alphacoronavirus) and 229E (Alphacoronavirus) circulate widely in the human population, causing mild respiratory illnesses like the common cold (*63*). Mouse Hepatitis Virus (MHV, Betacoronavirus lineage A) is a rodent pathogen and was used for resistance studies in Paxlovid licensing due to its harmlessness to humans (*76*, *77*). Lastly, poliovirus (Picornaviridae) is still causing sporadic local outbreaks, mostly through reverted, live attenuated vaccine strains and in unvaccinated populations (*78*, *79*).

We found that both nirmatrelvir and ensitrelvir strongly inhibited SARS-CoV-2 and SARS-CoV-1 main proteases. Nirmatrelvir had moderate activity against the main proteases of MERS-CoV and HKU9 and mild effects on NL63- and 229E-M^pro^. In a previous study, we also showed a weak effect of nirmatrelvir on MHV-M^pro^ by using Fluorescence Activated Cell Sorting (FACS) as more elaborate, but also more sensitive read-out method (*29*). Ensitrelvir however had only mild activity against the main proteases of MERS-CoV and HKU9.

To explore the potential relationship between sequence similarity among examined proteases and their response to inhibitors, we aligned the amino acid sequences of these proteases with that of SARS-CoV-2 as reference, given that the tested inhibitors were originally licensed for its main protease. Underscoring the different conservation degrees in distinct parts of the viral genome, the main proteases of SARS-CoV-2 and SARS-CoV-1 share high sequence similarity (96 %), despite these two viruses emerging 20 years apart. In contrast, spike proteins of SARS-CoV-2 Wuhan-1 and SARS-CoV-1 share only 76 % identity. Furthermore, the SARS-CoV-2 spike protein has acquired numerous mutations during the pandemic, whereas only one dominant M^pro^ mutation, P132H, appeared in the Omicron variant (*80*). Taken together, sequence similarity correlated with inhibitor susceptibility in most of the tested proteases. An exception was MHV-M^pro^, which has a sequence identity of 51 % to SARS-CoV-2, comparable to that of MERS-CoV (51%), but was less responsive to nirmatrelvir. Structural analysis indicated strong backbone conservation, suggesting that the poor response stemmed from finer structural residue arrangements. Docking experiments revealed a higher calculated inhibition constant of nirmatrelvir against MHV-M^pro^ compared to proteases with similar sequence identity. In conclusion, our findings emphasize the importance of employing different methods to assess the potential of protease inhibitors against current and future CoVs, and caution against relying on models alone.

As the focus of this study was the applicability of SARS-CoV-M^pro^ inhibitors to MERS-CoV-M^pro^, we investigated the structural similarities of those two main proteases in particular. Despite an amino acid sequence identity of only 51% (*81*), the superimposition of their X-ray crystal structures revealed substantial structural similarity, including the conservation of the active site. To our knowledge, no specific antiviral has been developed directly targeting MERS-CoV thus far (*21*). One of the most promising potential treatments seemed to be passive immunotherapy using convalescent plasma, which is not as practical as a small molecule and certainly not as scalable in production (*82*). The last years however have seen massive efforts to find therapies for SARS-CoV-2, resulting in a pool of antivirals that might also be viable for MERS-CoV. We therefore examined the activity and inhibition of both SARS-CoV-2 and MERS-CoV main proteases using seven previously described protease inhibitors: nirmatrelvir (*41*), ensitrelvir (*43*), GC376 (*40*), PF-00835231 (*42*), bofutrelvir (*45*), compound **19** (*44*) and boceprevir (*40*). As expected, all tested inhibitors were effective against SARS-CoV-2-M^pro^. The most effective compounds against MERS-CoV-M^pro^ were GC376, nirmatrelvir and PF-00835231. Interestingly, ensitrelvir showed a significantly weaker effect on MERS-CoV-M^pro^ relative to SARS-CoV-2-M^pro^.

When examining nirmatrelvir interaction with active sites of SARS-CoV-2 and MERS-CoV main proteases in 2D maps, we observed the substitution M49L, previously linked with selective ensitrelvir resistance (*83*, *84*), and N142 / 145C. In a previous study, we identified the SARS-CoV-2-M^pro^ mutant N142D in nirmatrelvir selection experiments (*48*). Furthermore, substitutions H164 / 167Q and R188 / 191K also resembled substitutions from our recent mutation studies involving nirmatrelvir and ensitrelvir (SARS-CoV-2-M^pro^ mutants H163Q, H164N and R188W) (*47*, *48*). When analyzing the catalytic site interactions with ensitrelvir, we observed the natural variation T25M in MERS-CoV-M^pro^. Our recent study emphasized the significance of position 25 as a key interaction residue, evidenced by the emergence of specific ensitrelvir mutants (T25A, T25N) in SARS-CoV-2-M^pro^ (*47*). Consequently, we cloned MERS-CoV-M^pro^ mutant M25T, expecting decreased resistance. Surprisingly, M25T not only failed to sensitize MERS-CoV-M^pro^ to ensitrelvir but significantly increased its resistance. In addition, M25T was more susceptible to nirmatrelvir than the wild-type, whereas introducing T25M into SARS-CoV-2-M^pro^ did not affect resistance.

To explore resistance mechanisms of position 25 mutants, we studied the structure of the substrate-binding sites, which include four distinct subsites (S1’, S1, S2 and S4) (*85*). Non-covalent ensitrelvir has a binding affinity similar to nirmatrelvir (*65*, *86*). However, it differs in its binding mode from covalent inhibitors like nirmatrelvir or GC376, as it additionally binds in the top of the S1’ pocket (*87*). The S1’ pocket of MERS-CoV-M^pro^ is smaller than that of SARS-CoV-2-M^pro^, potentially causing lower affinity of compounds extending into this site (*88*), as in the case of ensitrelvir. The upper part of the S1’ pocket comprises two loops, one with M25 (residues 21 – 26) and the other lining the opposite side of the S1’ pocket (residues 42 – 49). In SARS-CoV-2-M^pro^, T25 interacts with C22 and V42 or C44 or all three residues, depending on the respective crystal structure, thereby stabilizing loop conformations. Similarly, the MERS-CoV-M^pro^ M25T mutant, according to our modelling, enhances protein stability by an additional hydrogen bond of T25 with C44 and leads to a movement of L49 towards H41. Interestingly, when comparing experimentally determined MERS-CoV-M^pro^ apo and nirmatrelvir bound structures, we found the loop containing L49 is also situated closer to H41 in the nirmatrelvir bound state. The MERS-CoV-M^pro^ M25T therefore mimics this L49 to H41 movement according to MD simulations. On the one hand, this stabilizes the smaller S1’ pocket less compatible with ensitrelvir binding, while on the other hand, it stabilizes the nirmatrelvir bound conformation. Therefore, M25T has different effects on the two inhibitors ensitrelvir and nirmatrelvir.

Nirmatrelvir was effective against various viral proteases, and most importantly for this study, against MERS-CoV-M^pro^. The molecular mechanism of nirmatrelvir inhibiting SARS-CoV-2-M^pro^ involves a covalent reaction with the catalytically active cysteine C145 (*28*, *65*), the functional analogous cysteine in MERS-CoV-M^pro^ being C148. To simulate nirmatrelvir use and resistance development against MERS-CoV-M^pro^, we selected a pool of mutants and evaluated those mutations in proximity to the inhibitor binding site. S147Y showed strong resistance to inhibitors, prompting us to inspect the structural conditions of this mutation. Tyrosine, having a larger side chain than serine, occupies nirmatrelvir’s binding site leading to a clash with the inhibitor. Interestingly, the tyrosine side chain occupies the exact same location, where cyclic substituents of M^pro^ inhibitors mimic interactions with the P1 glutamine of the substrate, suggesting that this mutant sterically hinders proper positioning of these inhibitor substructures. This could explain the observed resistance to nirmatrelvir and also indicate the loss of efficacy for other tested inhibitors, which similarly contain components occupying the S1 pocket.

Counterintuitively, A171S showed an increased affinity to nirmatrelvir. Given that this mutant only had a mild phenotype, we speculate that it is either compensatory or augmenting to a second mutation we found in this sample, D289G, or that it sequesters nirmatrelvir in a less optimal binding pose. Mutants S142G and S142R exhibited slower kinetics, but also increased nirmatrelvir resistance. In S142G, the disruption of the hydrogen bond with Q299, crucial for dimerization, probably reduces binding affinity between the protomers due to greater movement in this region and a lower chance of attaining the correct conformation. Additionally, Δ stability values indicated reduced stability and Log10 K* scores decreased dimer affinity, which suggests a lower likelihood of adopting the nirmatrelvir binding conformation, providing a rationale for the observed nirmatrelvir resistance phenotype. The mutant S142R demonstrated stronger resistance to nirmatrelvir than S142G. Besides lacking the Q299 hydrogen bond, the larger size of arginine compared to serine may lead to steric clashes between the protomers. Supporting our hypothesis, S142R showed even lower stability than S142G and also reduced affinity.

Our study has limitations. First and foremost, we were unable to test inhibition and resistance phenotypes in authentic MERS-CoV. Although the tools described in this study were developed precisely to avoid working and performing gain-of-function research with this dangerous pathogen, confirming the results in MERS-CoV as proof-of-concept would have been useful. Consequently, lacking replicating MERS-CoV, animal experiments could also not be done. Furthermore, we did not test all inhibitors featured in this study against all available viral proteases.

In conclusion, our findings suggest that SARS-CoV-2-M^pro^ inhibitors, especially nirmatrelvir or its derivates, could be used to develop specific antiviral drugs against MERS-CoV-M^pro^. Furthermore, we pointed out the broad applicability of nirmatrelvir to different viral proteases and possible resistance pathways of MERS-CoV-M^pro^.

## MATERIALS AND METHODS

### Study Design

This study aimed to assess protease inhibitors, developed or already licensed against SARS-CoV-2, for their effectiveness against MERS-CoV-M^pro^. To elucidate specific resistance patterns of both main proteases in relation to ensitrelvir, we conducted directed mutagenesis. Additionally, nirmatrelvir resistant mutants were selected and characterized. We employed previously described VSV-based cellular On and Off assays (*30*) and a VSV mutation selection tool (*29*). Cross validation to confirm results and suggest mechanisms were performed with a biochemical assay and molecular modelling. Viral titers were determined using TCID_50_. Resistance measurement read-outs were obtained through fluorescence-based detection facilitated by a FluoroSpot reader. BHK-21 (hamster) and 293T (human) cells were used to study viral replication and resistance phenotypes. No other animal or human material was used to perform this study. Experiments were not blinded.

### Cloning strategies

To create a MERS-CoV-M^pro^ dependent VSV construct, the intergenic region between G and L of a VSV-Indiana plasmid (**plasmid S1**) was replaced by MERS-CoV-M^pro^, generating VSV-G-M^pro^-L / VSV-MERS-M^pro^ (**plasmid S2**). The VSV plasmid was digested with restriction enzymes KpnI and HpaI (New England Biolabs, NEB), excising a C-terminal part of G, the intergenic region and a N-terminal part of L. Insert PCRs were carried out to clone two MERS-CoV-M^pro^ variants with cleavage site lengths of seven and eight amino acids (7aa, 8aa). The first fragment included C-terminus of G with an additional overhang to the N-terminal cleavage site of M^pro^, amplified using primers G-33n-before-KpnI-for and G-rev. The M^pro^-fragments (7aa and 8aa) were amplified with primers G-cut1-*aa-for and cut2-*aa-L-rev. Finally, the missing N-terminal L sequence was amplified with primers L-for and L-33n-after-HpaI-rev. For subsequent Gibson assembly (NEB, USA), the fragments were joined in a fusion PCR using the outer primers G-33n-before-KpnI-for and L-33n-after-HpaI-rev with three fragments as templates.

For MERS-CoV-M^pro^-On assays, we modified a lentiviral plasmid (pLenti CMVie-IRES-BlastR, Addgene accession: #11963) by replacing the blasticidin resistance cassette, removed by MscI and NotI (NEB), with a hygromycin resistance cassette, yielding pLenti CMVie-IRES-HygroR (**plasmid S3**). MERS-CoV-M^pro^ was codon-optimized for mammalian expression, synthesized by Integrated DNA Technologies (USA) and inserted intramolecularly into the VSV P / phosphoprotein (P:M^pro^:P) as described before (*30*). To increase sensitivity, a N-terminal glutamine to asparagine mutation (QtoN) was applied to M^pro^ (*30*). Briefly, we conducted PCR amplification of three fragments (P-Nterm, M^pro^ and P-Cterm) with overlapping sequences for subsequent fusion PCR using primer pairs Hygro-P-for / P-GGSG-rev, MERS-On-N-term-QtoN-for / MERS-On-C-term-rev and GGSG-P-for / Hygro-P-rev. This fusion PCR product was then integrated into the NheI- and PacI-digested pLenti CMVie-IRES-HygroR through Gibson assembly, generating MERS-On-N-term-QtoN (**plasmid S4**). The SARS-2-N-term-QtoN construct was cloned by mutagenesis of On wt with primers SARS-2-On-N-term-QtoN-for and -rev. The other On plasmids expressing different proteases (as in **Figure 5**) were cloned analogously to MERS-On-N-term-QtoN (*30*), using *-On-N-term-QtoN-for and *-GGSG-On-rev primers (**table S3**) with codon-optimized M^pro^ sequences for mammalian expression.

To generate a MERS-CoV-M^pro^-Off plasmid, pLenti CMVie-IRES-BlastR was digested with NheI and PacI. The VSV L polymerase sequence was amplified from **plasmid S1** with primers Blasti-L-for / L-blasti-rev and inserted into pLenti CMVie-IRES-BlastR via Gibson assembly, generating VSV-L-BlastR (**plasmid S5**). Subsequently, VSV-L-BlastR was cleaved by HpaI, which removed segments of the lentiviral vector and a small N-terminal part of L. The missing parts of the lentiviral vector and GFP were replaced through PCRs with primers Blasti-for / GFP-rev with SARS-2-M^pro^-Off (*30*) (GenBank accession number ON262564) as template. MERS-CoV-M^pro^ was amplified with primers GFP-cut1-MERS-Off-7aa-for / 7aa-MERS-Off-cut2-L-rev from MERS-M^pro^-On (*30*) and N-terminal L with primers L-33n-after-HpaI-rev / L-for from **plasmid S5**. These three fragments were merged via fusion PCR and cloned into VSV-L-BlastR digested with HpaI, generating MERS-M^pro^-Off (**plasmid S6**).

MERS-CoV-M^pro^-On and -Off point mutants were cloned by directed mutagenesis applied on the parental plasmids, using specific mutation primers (**table S3**). To amplify large plasmids, the Herculase II polymerase (Agilent, USA) was employed, with primer temperatures determined by the NEB primer temperature calculation website (https://tmcalculator.neb.com/#!/main) for each respective primer pair. The PCR elongation times were 10 minutes for 45 cycles. All cloning primers used in this study are listed in **table S3.**

### Cell lines

BHK-21 cells (American Type Culture Collection, ATCC) were cultured in Glasgow Minimum Essential Medium (GMEM) (Lonza) supplemented with 10 % fetal calf serum (FCS), 5 % tryptose phosphate broth, and 100 units/ml penicillin plus 0.1 mg/ml streptomycin (P/S) (Gibco). For the Human Embryonic Kidney (HEK) cell line 293T (293tsA1609neo, ATCC), Dulbecco’s Modified Eagle Medium (DMEM) supplemented with 10 % FCS, 2 % glutamine, 1 % P/S, 1x sodium pyruvate, and 1x non-essential amino acids (Gibco) was used.

### Virus recovery

VSV-MERS-M^pro^ was rescued in 293T cells as previously described (*29*, *89*). In brief, 293T cells were seeded a day prior to transfection and whole-genome VSV plasmids along with a helper plasmid mix containing T7-polymerase, N-, P-, G- and L expression plasmids were transfected into cells using CaPO_4_ (so-called “rescue”). We used 10 µM chloroquine to prevent lysosomal DNA degradation. After 6 to 16 hours, chloroquine was removed and cells were cultured until cytopathic effects occurred. Both 7aa and 8aa viruses were then passaged and plaque purified on BHK-21 cells, showing full replication capability. We used VSV-ΔP-dsRed for M^pro^-On and VSV-ΔL-dsRed for M^pro^-Off (*90*). The ΔP and ΔL viruses were cultivated on replication supporting 293T cells expressing VSV-P or VSV-L, respectively.

### Dose-responses and TCID_50_ assays

For dose-response experiments, 2 x 10^4^ BHK-21 cells per well were seeded in a 48-well plate one day before infection. Subsequently, plates were infected with nirmatrelvir and VSV-MERS-M^pro^ virus at a multiplicity of infection (MOI) of 0.01. Nirmatrelvir concentrations started at 100 µM and were diluted 1:3 eleven times. One row was left without nirmatrelvir. Each condition had three replicates. After 24 hours, supernatants were collected and titrated. After six days, titers were calculated based on the Kaerber method (*91*) to determine the 50 % tissue culture infective dose (TCID_50_).

### Cellular M^pro^-On screening assay

For the M^pro^-On construct, two vectors were used to trans- and infect 293T cells: a plasmid and a VSV replicon, respectively. First, an intramolecular insertion of the MERS-CoV main protease sequence into the VSV P protein (P:M^pro^:P) was carried out to introduce MERS-CoV-M^pro^ into an established reporter system (*38*). The VSV replicon (VSV-ΔP-dsRed) comprised VSV leader / trailer sequences, nucleoprotein (N), a dsRed reporter gene, phosphoprotein (P), matrix protein (M), glykoprotein (G) and polymerase (L) (*31*). Co-expression of these two vectors in the same cells led to autocatalytic cleavage of the P protein by M^pro^, impeding virus replication. Adding an inhibitor restored P protein function, enabling virus replication and dsRed signal accumulation (“gain-of-signal”). We used the N-terminal QtoN version of this assay (*30*), which increased sensitivity. M^pro^-On assays were used to evaluate protease inhibitors, initially developed for SARS-CoV-2 treatment, against MERS-CoV-M^pro^ and other viral proteases. Furthermore, we used this assay to study resistance in “MERS-CoV to SARS-CoV-2” mutants. For the assay, 3 x 10^5^ 293T cells per well were seeded in 6-well plates and transfected with VSV-P:M^pro^:P plasmids a day later, using TransIT-^pro^ (Mirus Bio LLC). After 10 hours of transfection, cells were trypsinized and seeded in 96-well plates at 2 x 10^4^ cells / well in 50 µl complete growth medium. Then, inhibitor dilutions and VSV-ΔP-dsRed virus (MOI: 0.1) were added in 50 µl complete growth medium to wells. 48 hours later, fluorescence signals were measured using a FluoroSpot reader and graphs compiled with GraphPad Prism 10 (GraphPad Software, Inc.).

### M^pro^-Off screening assay for resistance mutations

The MERS-CoV-M^pro^-Off assay was based on replication-incompetent VSV-ΔL-dsRed replicons containing a dsRed reporter gene (*30*). Off-plasmids were generated by tagging the N-terminal VSV L polymerase with M^pro^, eGFP and a protease cleavage site (GFP-M^pro^-L). VSV-ΔL-dsRed replication was dependent on the autocatalytic processing by M^pro^, releasing a functional L polymerase for replication. Introducing an inhibitor blocked cleavage and suppressed the accumulation of dsRed signal (“loss-of-signal”). Off assays were conducted to confirm M^pro^-On assays and to assess resistance of SARS-CoV-2 and MERS-CoV main protease mutants. 3 x 10^5^ 293T cells per well were seeded in 6-well plates and transfected one day later with GFP-M^pro^-L plasmids using TransIT-^pro^ (Mirus Bio LLC). After 10 hours, cells were trypsinized and seeded into 96-well plates with 2 x 10^4^ cells / well in 50 µl complete growth medium. Subsequent addition of an inhibitor in 50 µl complete growth medium, followed by infection with VSV-ΔL-dsRed virus particles (MOI: 0.1), yielded dsRed-positive cells. Fluorescent spots were quantified every 12 hours by a FluoroSpot counter within the first four days after infection.

### M^pro^-Off replication kinetics

To assess SARS-CoV-2 and MERS-CoV main protease kinetics along with mutants thereof, we adapted the M^pro^-Off assay for multiple measurements. We used a phenole-red free medium (FluoroBrite^TM^) to reduce background fluorescence by the medium. Eight replicates per M^pro^ or mutant were infected with VSV-ΔL-dsRed at an MOI of 0.1 like in M^pro^-Off assays, but without inhibitor. Measurements were taken every 12 hours and graphs compiled with GraphPad Prism 10. We quantified and compared protease kinetics using “TM_50_”, denoting the time at which the signal reached its half-maximum, in analogy to IC_50_ values.

### FluoroSpot read-out

Fluorescence signals in M^pro^-On and -Off assays were assessed using a FluoroSpot/ELISpot/ImmunoSpot counter (CTL Europe GmbH, Bonn, Germany). Concentrical scanning was performed with CTL switchboard 2.7.2. A 570 nm laser was used for dsRed excitation, and fluorescence was captured using the D_F_R triple-band filter. In the presence of FluoroBrite^TM^ medium, we set the counting area at 70 %, corresponding to the area that actually contains cells and signal. Without medium, 80 % of the well area was counted to minimize well boundary reflections. Automatic fiber exclusion was applied during scanning. To ensure comparability between M^pro^-On and -Off signals, we normalized dsRed events with two strategies. In M^pro^-On, the highest compound concentrations could not yield equivalent values due to distinct responses among mutants. Thus, normalization was performed against the highest mean value within the experiment. In M^pro^-Off, all constructs achieved the same signal intensity and variation depended mostly on technical parameters such as cell count. Thus, we normalized the signal based on the individual highest mean value for each construct. Neither normalization affected IC_50_ or TM_50_ values, but improved visualization. In both assays, fluorescent spot counts (y-axis) were plotted against protease inhibitor concentration (x-axis). Subsequent analysis involved extrapolating IC_50_ values, as described in the **Graph generation, IC_50_ and TM_50_ calculations using GraphPad** section.

### Mutation selection assay

For the selection of mutants, we only used the 7aa virus variant due to its analogous dose-response curve with the 8aa variant and our previous extensive experience with SARS-CoV-2-M^pro^ containing 7aa cleavage sites. We seeded 10,000 BHK-21 cells per well in 48 wells of a 96-well plate one day before infection. VSV-MERS-M^pro^ was added at a MOI of 0.01, along with 40 µM nirmatrelvir. Wells exhibiting cytopathic effects within 48-72 hours were further passaged. Nirmatrelvir concentrations were increased in each passage (1^st^: 40, 2^nd^: 50, 3^rd^: 60, 4^th^: 70 and 5^th^: 80 µM) over five consecutive resistance passages. In the final step, supernatants from cytopathic wells were collected and viral RNA was isolated through a semiautomated RNA extraction instrument called Easymag (BioMerieux^TM^, France). RNA purification was followed by cDNA synthesis with RevertAid RT Reverse Transcription Kit (Thermo Fisher Scientific). Sequencing was conducted via Nanopore or Sanger (MicroSynth AG) methods. Obtained mutations were mapped to reference MERS-CoV-M^pro^ sequences using Geneious Prime 2023.0.1 and were manually documented in Excel spreadsheets. Silent mutations and known sequencing artefacts (such as miscounts of stretches of the same nucleotide) were excluded from further study.

### Nanopore sequencing workflow

To create libraries for sequencing, we used the Rapid Barcoding Kit SQK-RBK110.96 (Oxford Nanopore Technologies, ONT) and multiplexed up to 96 samples for sequencing with R9.4.1 flowcells on the MinION Mk1B platform (ONT). Raw signal data, stored in pod5 files, were converted to nucleotide sequences and saved in fastq files, using the super high accuracy model of Guppy (version 6.5.7, ONT), which also performed demultiplexing, as well as trimming of adapter and barcode sequences. Next, reads falling outside the range of 200 to 1800 bp, and with an average PHRED scale quality score below Q15, were removed with SeqKit (version 2.4.0) (*92*). After filtering, reads were aligned to the reference sequence in minimap2 (version 2.22) (*93*), using the preset for Oxford Nanopore (map-ont), and then sorted and indexed with SAMtools (version 1.13) (*94*). To enable checking for sufficient depth, the read depth at each position was computed using the samtools depth command. Calling of single nucleotide variants was facilitated by running LoFreq (v2.1.3.1) with the --min-cov parameter set to 300) (*95*). Finally, the resultant VCF files were loaded in Geneious Prime 2023.0.1 and the called mutations manually examined for plausibility.

### Graph generation, IC_50_ and TM_50_ calculations using GraphPad

IC_50_ calculations and statistical analysis for all assays were conducted using GraphPad Prism 10. IC_50_ values were calculated using pre-set sigmoidal models from GraphPad, specifically the 4-parameter logistic (4PL) model with concentration (X) as the variable.

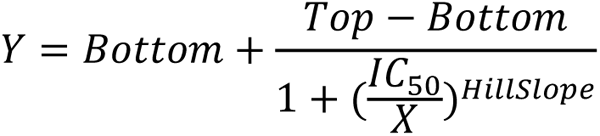

Under certain conditions, our assays exhibited slightly bell-shaped curves due to signal fluctuations before (Off) or after (On) the signal plateau. The initially mildly lower signal and then increase in the Off assay might result from rapid cell death without an inhibitor, while slower cell death at low inhibitor doses allowed more cell division, generating more “substrate” (cells and virus) and increased dsRed production. In the On assay, the decrease after the plateau can be attributed to inhibitor toxicity at high concentrations. These bell-shape curves could potentially lead to overfitting issues with the sigmoidal models, resulting in overly steep decreasing or increasing curves. Although this generally did not significantly affect IC_50_ values, we have taken measures to address these unrealistically steep slopes. To that end, we constrained the HillSlope parameter to 2 (positive slope) for the On assay and -3 (negative slope) for the Off assay. We defined TM_50_ as proxy for M^pro^ kinetics using the same formula as for IC_50_, but with hours as input instead of inhibitor concentrations and without additional slope constrains.

### Structure alignment and RMSD calculation

**Table S1** shows a superposition matrix generated in ICM genome browser 3.9-3a, using residues 9 – 194 of chains A (inhibitor binding domain) from structures of different viruses. The function Secondary Structure Matching (SSM) superposition (*96*) was applied on Cα backbone atoms to retrieve RMSD in Ångström (Å). MERS-CoV-M^pro^ (PDB ID: 5C3N) and SARS-CoV-2-M^pro^ (PDB ID: 7VH8) were aligned using ChimeraX matchmaker with the Needleman-Wunsch algorithm. RMSD between 253 pruned atom pairs was 0.906 Å and across all 297 pairs 1.502 Å.

### Inhibitor interaction with SARS-CoV-2 and MERS-CoV main proteases

To visualize the interactions between protease inhibitors and main proteases of SARS-CoV-2 and MERS-CoV in 2D representations, we used Maestro (Schrödinger Inc., USA). As input, we used 3D inhibitor-bound monomer protease structures from the PDB. 2D images depicting M^pro^ active sites with nirmatrelvir and ensitrelvir were put out by the software. An ensitrelvir-bound MERS-CoV-M^pro^ structure was not available, we generated the complex structure with Autodock 4.

### Molecular docking with AutoDock 4

Given the small conformational variations between the different proteases, even at the level of side-chain torsion angles, we chose a docking model that uses a rigid protein but a flexible linker. For ensitrelvir, we employed AutoDock 4 and AutoDock Tools (*97*) for rigid docking. All protease structures were superimposed so that they share the same grid. The following template structures from the PDB (https://www.rcsb.org/) were used: SARS-CoV-2 (PDB ID: 8DZ1), SARS-CoV-1 (PDB ID: 7ZQW), MERS-CoV (PDB ID: 7VTC), HKU9 (SWISS Model), NL63 (PDB ID: 5GWY), 229E (PDB ID: 2ZU2), MHV (PDB ID: 6JIJ) and Polio (PDB ID: 4DCD). For ensitrelvir, the 3D conformer SDF file (PubChem CID: 162533924) was downloaded from the PubChem database (https://pubchem.ncbi.nlm.nih.gov/). The search space of the pocket was defined as a 40 Ångström box centered on the thiol group of cysteine 145 (SARS-CoV-2 numbering). For each protease structure, the flexible ligand was subjected to 500 docking runs, and the best hits were clustered based on the highest negative free energy (ΔG) calculated by AutoDock 4 and converted into Kd values using the equations ΔG° = -RT ln Ki and Kd = 1/Ki. For covalent docking of nirmatrelvir, the same methods were used as recently reported (*98*) with the same template files from the PDB as described for ensitrelvir. The 3D conformer SDF file for nirmatrelvir (PubChem CID: 155903259) was retrieved from PubChem. The search space of the pocket, number of runs and free energy-based clustering were used as described above for ensitrelvir.

### Stability and affinity predictions and covalent docking with Schrödinger software packages

The PDB structures 5C3N (*24*) (MERS-CoV-M^pro^ Apo), 7VTC (*49*) (MERS-CoV-M^pro^ bound to nirmatrelvir), and 7VLQ (*49*) (SARS-CoV-2-M^pro^ bound to nirmatrelvir) were used in this study. All structures underwent preparation using the Protein Preparation Wizard tool in Maestro Schrödinger version 2022-3 (*99*, *100*), with default settings applied except for converting seleno-methionines to methionines. Depending on the experiment, water molecules were deleted or kept. All stability predictions were calculated with the residue_scanning_backend.py module of BioLuminate by Schrödinger (*55–58*), using default settings as previously described (*48*). Osprey version 3.3 (*59–62*) was employed for affinity predictions between both protomers, as previously described (*48*). Briefly, Osprey calculated Log_10_ K* scores, estimating the binding affinity. YAML files were created as detailed in the Guerin et al. STAR Protocol (*101*). For dimer affinity calculations, chain B was considered as the ligand. The stability threshold was disabled, epsilon defined as 0.03, and mutant side chain conformations were computed as continuously flexible. Osprey is an open-source software accessible for free at https://github.com/donaldlab/OSPREY3. Covalent docking was performed with Maestro version 2022-3 (*51–54*). C148 was picked as reactive residue, with the reaction type defined as “Nucleophilic Addition to a Triple Bond”. MM-GBSA scoring was enabled and up to 10 output poses were retrieved. Before docking, nirmatrelvir was prepared using the default settings of the LigPrep module in Maestro (Schrödinger Release 2022-3: LigPrep, Schrödinger, LLC, New York, NY, 2022).

### Molecular dynamic simulation and data analysis with GROMACS software packages

Molecular dynamic (MD) simulations were conducted in GROMACS version 2022.4 (*102–105*) using the Charmm36 (version July 2022) (*106*) force field for topology preparation (*107*) and calculating the parameterization of proteins, solvents and ions. The calculation was done in a cuboid with at least 10 Å distance to the box boundaries. The system was solvated with the SPC/E (*107*, *108*) water system and the net charge was neutralized with sodium ions. The system underwent minimization and equilibration processes, including 100 ps runs using the NVT and NPT ensemble, where the atom count N of the systems is 83142 (5C3N wt (*24*)) and 83145 (5C3N M25T). During minimization, the system was set at 1 bar pressure and 300 K temperature. Each production run spanned 100 ns with time steps of 2 fs and was repeated twice, resulting in three production runs for each protein. Output frames were saved every 10 ps. Temperature was maintained at 300 K and pressure at 1 bar, using the Berendsen thermostat (*108*) and the C-rescale pressure coupling (*108*, *109*). Bond parameters were defined by the LINCS (*110*) algorithm, and long-range electrostatics by the Particle-Mesh Ewald algorithms (*111*). Trajectory analysis was conducted by the GROMACS analysis tool and plots were generated in Grace Version 5.1.25 (https://plasma-gate.weizmann.ac.il/Grace/).

### Additional software

Mutated residues were highlighted in the 3D structure of MERS-CoV-M^pro^ using ChimeraX. Heat maps were generated in GraphPad. Figures were compiled with Inkscape version 1.1.

### Protein expression and purification

M^pro^ with native termini was expressed using the CASPON platform technology (*112*). Briefly, M^pro^ with an N-terminal CASPON tag (CASPON-M^pro^) was expressed in *E.coli* followed by purification with Ni-NTA affinity chromatography and tag removal with the CASPON™ enzyme 1.0 provided by Boehringer Ingelheim RCV GmbH (Germany). Wild-type CASPON-M^pro^ construct was codon-optimized for *E. coli* expression and ordered from GeneArt / Thermo Scientific (USA) (**plasmid S7**).

Point mutations to generate the mutant variants were introduced using the Q5 polymerase from New England Biolabs (NEB, USA). Primers M^pro^-expr.-S147Y-for and –rev (**table S3**) for point mutagenesis were acquired from Sigma Aldrich / Merck (Germany). Expression and purification of constructs were conducted as previously described (*48*).

### Hexahistidine tag removal by negative IMAC

For tag removal, CASPON-M^pro^ was incubated with CASPON™ enzyme 1.0 in a ratio of 1:25 (m:m) at 25°C for 15 minutes. Native tag-free M^pro^ was then obtained by negative IMAC purification with a manually packed Ni-NTA column. The flowthrough yielded M^pro^ with native N- and C-termini, as both the tag and CASPON™ enzyme 1.0 were bound to the Ni-NTA resin. Imidazole and NaCl present in the Ni-NTA running buffer were eliminated by using a HiTrap Desalting 5 mL (GE Healthcare) column.

### Cross-validation with biochemical M^pro^ inhibition assay

Confirmation of the resistance phenotype of mutant S147Y was performed with a biochemical assay based on a BPS Biosciences kit (catalogue number #78042-2; USA). The SARS-CoV-2-M^pro^ of the kit was replaced by an in-house produced MERS-CoV-M^pro^ and a mutant thereof (S147Y), as described previously (*48*). Three hundred ng per reaction of the main proteases were prepared in 30 µl assay buffer (20 mM Tris/HCl pH = 8, 150 mM NaCl, bovine serum albumin 0.1 mg/ml, 1 mM dithiothreitol – DTT). A three-fold serial dilution of nirmatrelvir was added in 10 µl assay buffer and incubated for 30 minutes at 37°C. Then, 10 µl of the fluorogenic substrate Ac-Abu-Tle-Leu-Gln-MCA (Peptide Institute, Japan) were added and incubated over night at room temperature. Fluorescence was measured by excitation at 365 nm and read-out at 415 – 445 nm emission with a Glomax Explorer fluorometer (Promega, USA).

### Data exclusion

When generating dose-response curves, an outlier data point from VSV-MERS-M^pro^-8aa at 100 µM nirmatrelvir, which was 10,000-fold higher than the others, was identified to be a mutant by sequencing and consequently excluded from the graph. Automated data exclusion by the FluoroSpot software removed large autofluorescent particles like fibers, hairs and other contaminants. Additionally, manual particle exclusion, as recommended by the manufacturer, was performed for quality control. Nanopore sequencing artefacts, like miscounted homomeric stretches of nucleotides, were also excluded from further analysis.

## Supporting information

Alignments_S1-4

Sequences_S1-8

## ACKNOWLEDGEMENTS

We want to thank Seyed Arad Moghadasi, PhD from the Howard Hughes Medical Institute of Prof. Reuben Harris for useful discussions, insights and providing different protease expression plasmids. We also want to thank Assoc. Prof. Jens Carlsson from the Science for Life Laboratory at the Uppsala University for providing compound **19**. Furthermore, the authors thank Boehringer Ingelheim RCV GmbH & Co KG (Austria) for providing CASPON™ enzyme 1.0 and for general support on application of CASPON™ technology.

## Author contributions

E.H. conceived the initial concept. E.H., L.K, S.R. and F.C. designed the experiments. E.H. conceived the cloning strategies. E.H. and S.R. generated recombinant viruses. E.H., H.S., L.K., S.R., B.S and F.C. performed experiments. B.S produced recombinant proteases. H.S. and T.K. performed molecular modelling. E.H. and L.K. wrote the manuscript. D.v.L. and E.H. provided the infrastructure and funding to the project. D.B. sequenced VSV-M^pro^ mutants. All authors read and approved the final manuscript.

## Data and materials availability

All pertinent data to support this study are included in the manuscript and supplementary files. If required, further data supporting the findings are available upon request from: (https://www.linkedin.com/in/emmanuel-heilmann-910208218/) (https://www.linkedin.com/in/francesco-costacurta/) (https://www.i-med.ac.at/virologie/)

