## Supplements for "Identification of key residues in MERS-CoV and SARS-CoV-2 main proteases for resistance against clinically applied inhibitors nirmatrelvir and ensitrelvir"

**This PDF file includes:**

Fig. S1 to S6
Tables S1 to S3
Captions for alignments S1 to S4
Captions for plasmids S1 to S7
Captions for sequences S1 to S8

**Other Supplementary Materials for this manuscript include the following:**

Alignments S1 to S4
Plasmids S1 to S7
Sequences S1 to S8

### Supplementary material

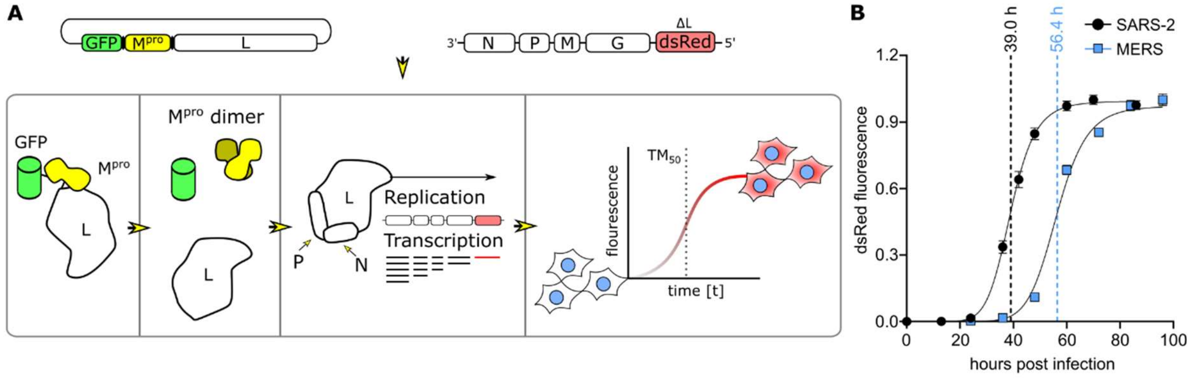

**Figure S1. Adaptation of the M$^{pro}$-Off assay for replication kinetics.** (**A**) The M$^{pro}$-Off assay was adapted by using GFP-M$^{pro}$-L and VSV-ΔL-dsRed without applying a protease inhibitor. Fluorescence signals are measured in regular intervals and plotted against time. $TM_{50}$ indicates the time required for the curve to reach half of its maximum value at which the signal plateaus. (**B**) M$^{pro}$-Off replication kinetics of SARS-CoV-2-M$^{pro}$ wt and MERS-CoV-M$^{pro}$ wt. Data are depicted as means of n = 8 biologically independent replicates per condition. Dotted lines represent the $TM_{50}$ value (hours post infection, hpi). At 80 hpi, both SARS-CoV-2 and MERS-CoV main proteases exhibit a plateauing dsRed signal.

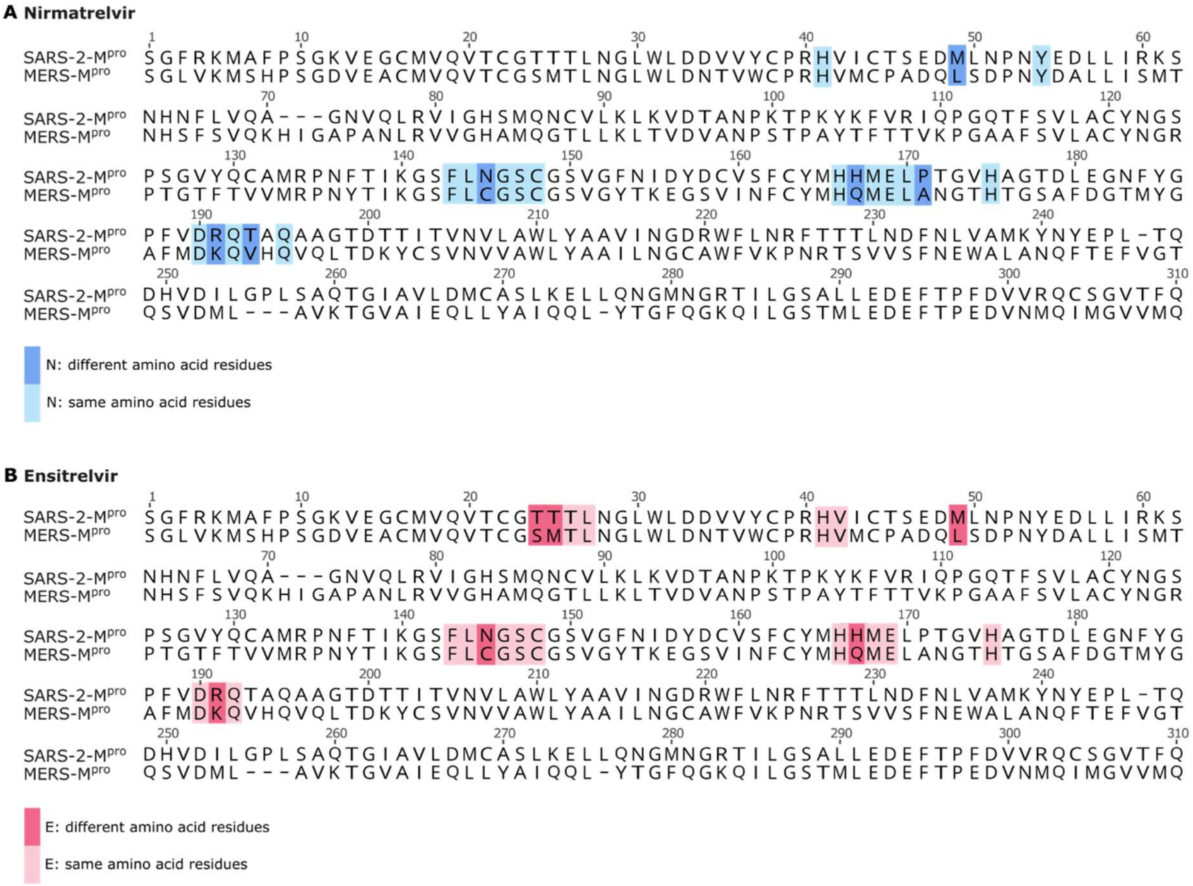

**Figure S2. Sequence alignments of SARS-CoV-2 and MERS-CoV main proteases highlighting the amino acid residues involved in inhibitor interaction.** (**A**) Natural variation of amino acid residues (dark blue) and same residues (light blue) that interact with nirmatrelvir. (**B**) Natural variation in amino acid residues (pink) and same residues (light pink) that interact with ensitrelvir. Interacting residues located within 4 Å of the inhibitor are included.

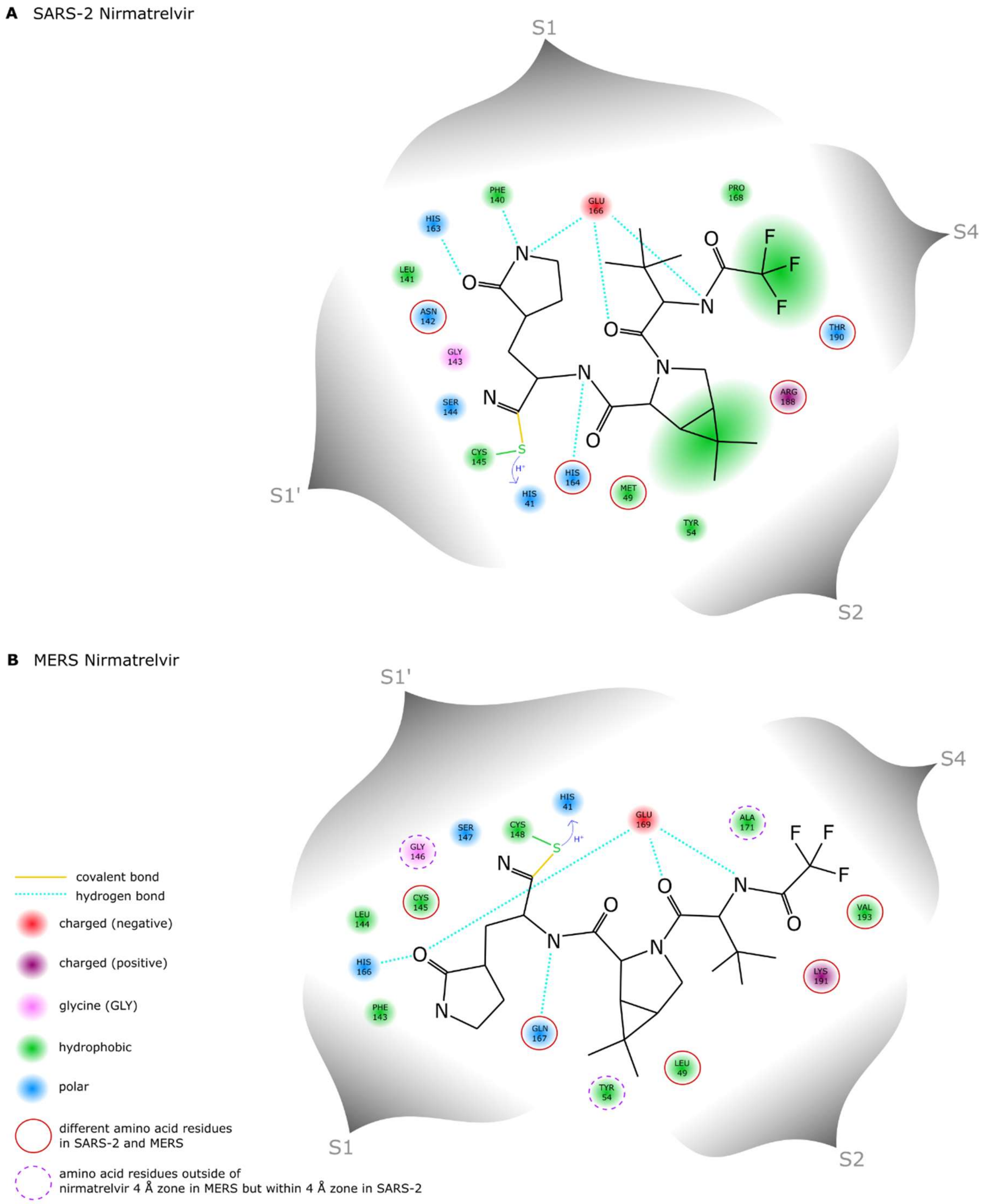

**Figure S3. Nirmatrelvir 2D binding maps with the catalytic sites of SARS-CoV-2 and MERS-CoV main proteases.** (**A**) The molecular mechanism of nirmatrelvir inhibiting SARS-CoV-2-M$^{pro}$ involves a reversible covalent reaction with the active site cysteine (C145). The nitrile group of nirmatrelvir serves as an electrophilic warhead, forming a covalent bond (yellow line) with the nucleophilic thiolate group of the catalytically active C145, situated within the S1' subsite. Residues 141 - 145 form an oxyanion loop harboring the C145 - H41 catalytic dyad. Histidine (H41) plays a crucial role in deprotonating (blue $H^+$) the C145 sulfur atom (green S), while G143 and C145 together create an oxyanion hole through their main chain amide NHs. Within the S1 interaction site, nirmatrelvir forms a hydrogen bond (turquoise dotted line) with H163 and three hydrogen bonds with E166. In the S2 subsite, nirmatrelvir's dimethylcyclopropyl proline (DMCP) is surrounded by hydrophobic interactions (green spheres). Similar interactions exist to stabilize the S4 pocket. (**B**) When examining the MERS-CoV-M$^{pro}$ catalytic site interacting with nirmatrelvir, some amino acid residues (red circles) differ from SARS-CoV-2-M$^{pro}$ interaction. There are the substitutions M49L, H164 / 167Q and R188 / 191K. Furthermore, the amino acid stretch 141 - 145, forming an oxyanion loop in SARS-CoV-2-M$^{pro}$, harbors the substitution N142 / 145C in MERS-CoV-M$^{pro}$, potentially reducing polar interactions. The amino acid variation T190 / 193V exchanges a polar with an apolar residue. G143 in SARS-CoV-2-M$^{pro}$ is essential for building a stabilizing oxyanion hole. The analogous glycine in MERS-CoV-M$^{pro}$ is outside of the nirmatrelvir 4 Å zone (purple dashed circle). The biochemical properties of interacting residues are displayed in red (negatively charged), purple (positively charged), green (hydrophobic) and blue (polar). Different binding types are also listed in the legend. Amino acid abbreviations: ALA (A), ARG (R), ASN (N), CYS (C), GLN (Q), GLU (E), GLY (G), HIS (H), LEU (L), LYS (K), MET (M), PHE (F), PRO (P), SER (S), THR (T), TYR (Y), VAL (V). 2D maps were generated with Inkscape, focusing on the interacting residues located within 4 Å from nirmatrelvir.

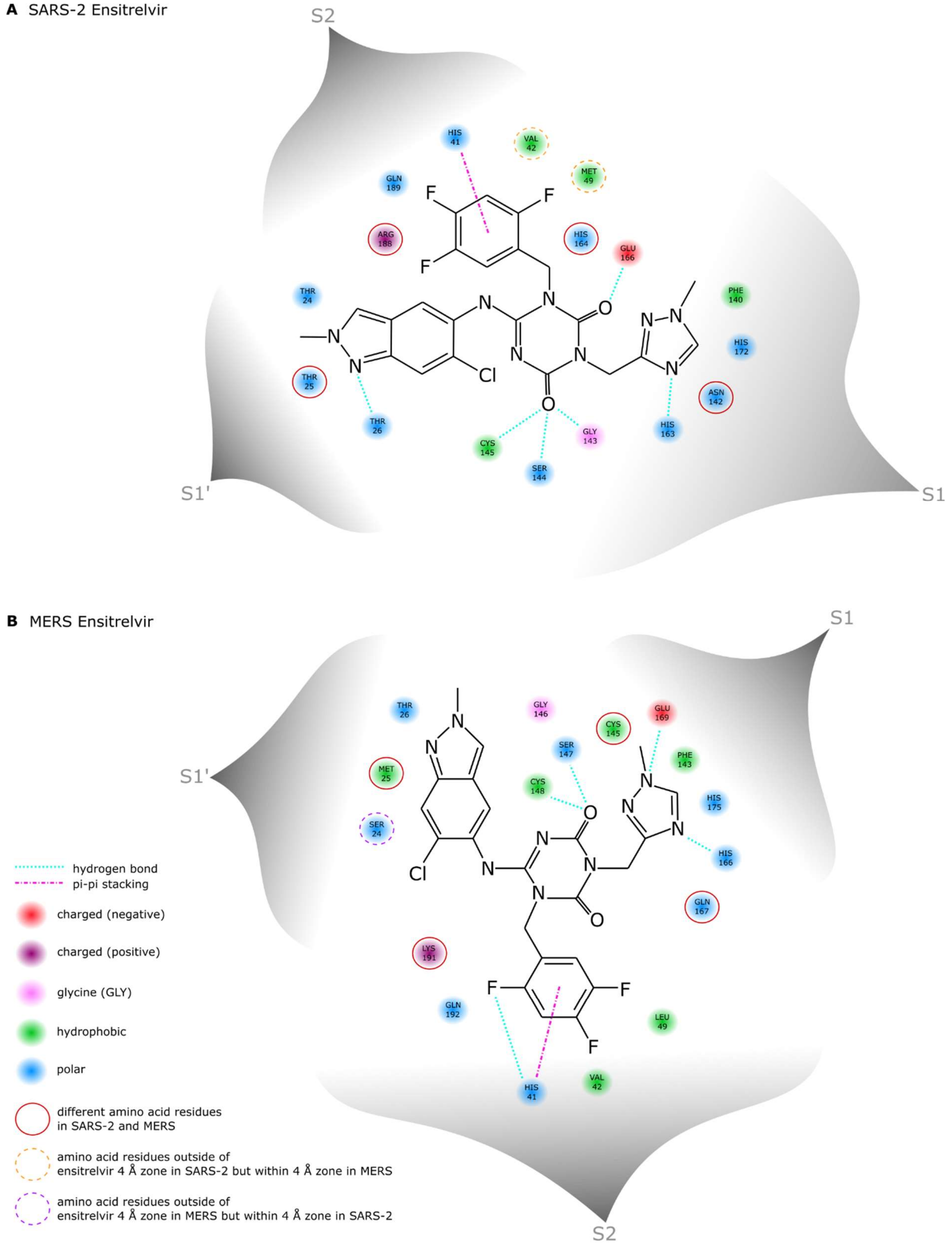

**Figure S4. Ensitrelvir 2D binding maps with the catalytic sites of SARS-CoV-2 and MERS-CoV main proteases.** (**A**) Ensitrelvir predominantly interacts with the S1', S1 and S2 subsites of SARS-CoV-2-M$^{pro}$, with fewer interactions within S4. The substrate pocket S1' harbors ensitrelvir's 6-chloro-2-methyl-2H-indazole moiety, which interacts with T26 through a hydrogen bond (turquoise dotted line). The interaction network within the S1 subsite consists mainly of stabilizing hydrogen bonds with E166, F140 and H172. H163 forms a hydrogen bond with the 1-methyl-1H-1,2,4-triazole group of ensitrelvir. In the S2 subsite, the 2,4,5-

trifluoromethyl of ensitrelvir engages in a pi-pi stacking interaction (pink dashed line) with the sidechain of H41. C145, G143 and Q189 are involved in the hydrogen bonding network stabilizing ensitrelvir. (**B**) Comparing the catalytic site residues interacting with ensitrelvir in MERS-CoV-$M^{pro}$, there are natural variations (red circles) from SARS-CoV-2-$M^{pro}$. In subsite S1', there are the substitutions T24S and T25M. S24 is not within the ensitrelvir 4 Å zone in MERS-CoV-$M^{pro}$ (purple dashed circle). M25 in MERS-CoV-$M^{pro}$ has a hydrophobic side chain, T25 in SARS-CoV-2-$M^{pro}$ a polar side chain. There are the substitutions N142 / 145C, H164 / 167Q and R188 / 191K. M49L is another natural variation, exchanging the hydrophobic amino acid residue of methionine (yellow dashed circle) outside of ensitrelvir's 4 Å zone with a bulkier, positively charged residue of lysine. The biochemical properties of interacting residues are displayed in red (negatively charged), purple (positively charged), green (hydrophobic) and blue (polar). Different binding types are also listed in the legend. Amino acid abbreviations: ARG (R), ASN (N), CYS (C), GLN (Q), GLU (E), GLY (G), HIS (H), LEU (L), LYS (K), MET (M), PHE (F), SER (S), THR (T), VAL (V). 2D maps were generated with Inkscape, focusing on the interacting residues located within 4 Å from ensitrelvir.

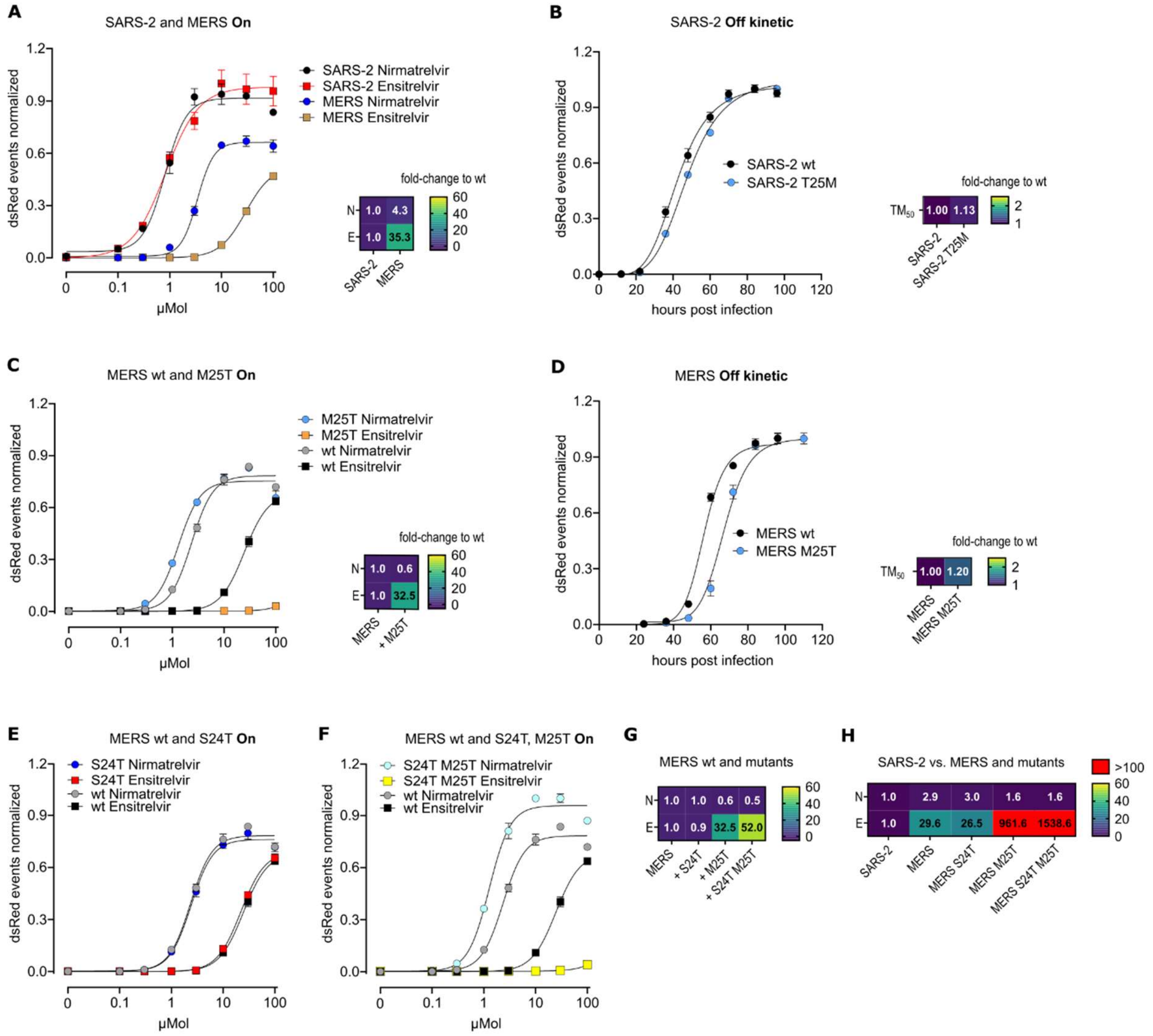

**Figure S5. Assessing nirmatrelvir (N) and ensitrelvir (E) efficacy against "MERS-CoV to SARS-CoV-2" mutants.** (**A**) $M^{pro}$-On assays of SARS-CoV-2-$M^{pro}$ wt and MERS-CoV-$M^{pro}$ wt with heat map showing $IC_{50}$ fold changes. (**B**) $M^{pro}$-Off replication kinetics of SARS-CoV-2-$M^{pro}$ wt and mutant T25M with heat map showing $TM_{50}$ fold changes. (**C**) $M^{pro}$-On assays of MERS-CoV-$M^{pro}$ wt and mutant M25T with heat map showing $IC_{50}$ fold changes. (**D**) $M^{pro}$-Off replication kinetics of MERS-CoV-$M^{pro}$ wt and mutant M25T with heat map showing $TM_{50}$ fold changes. $M^{pro}$-On assays of MERS-CoV-$M^{pro}$ wt and mutant S24T (**E**) and double mutant S24T / M25T (**F**). (**G**) Heat map showing $IC_{50}$ fold changes of (**E**), (**C**) and (**F**) compared to MERS-CoV-$M^{pro}$ wt. (**H**) Heat map showing $IC_{50}$ fold changes of (**A**), (**E**), (**C**) and (**F**) compared to SARS-CoV-2-$M^{pro}$ wt. $M^{pro}$-On assays are n = 3 biological replicates per condition. $M^{pro}$-Off kinetics are n = 8 biological replicates per condition.

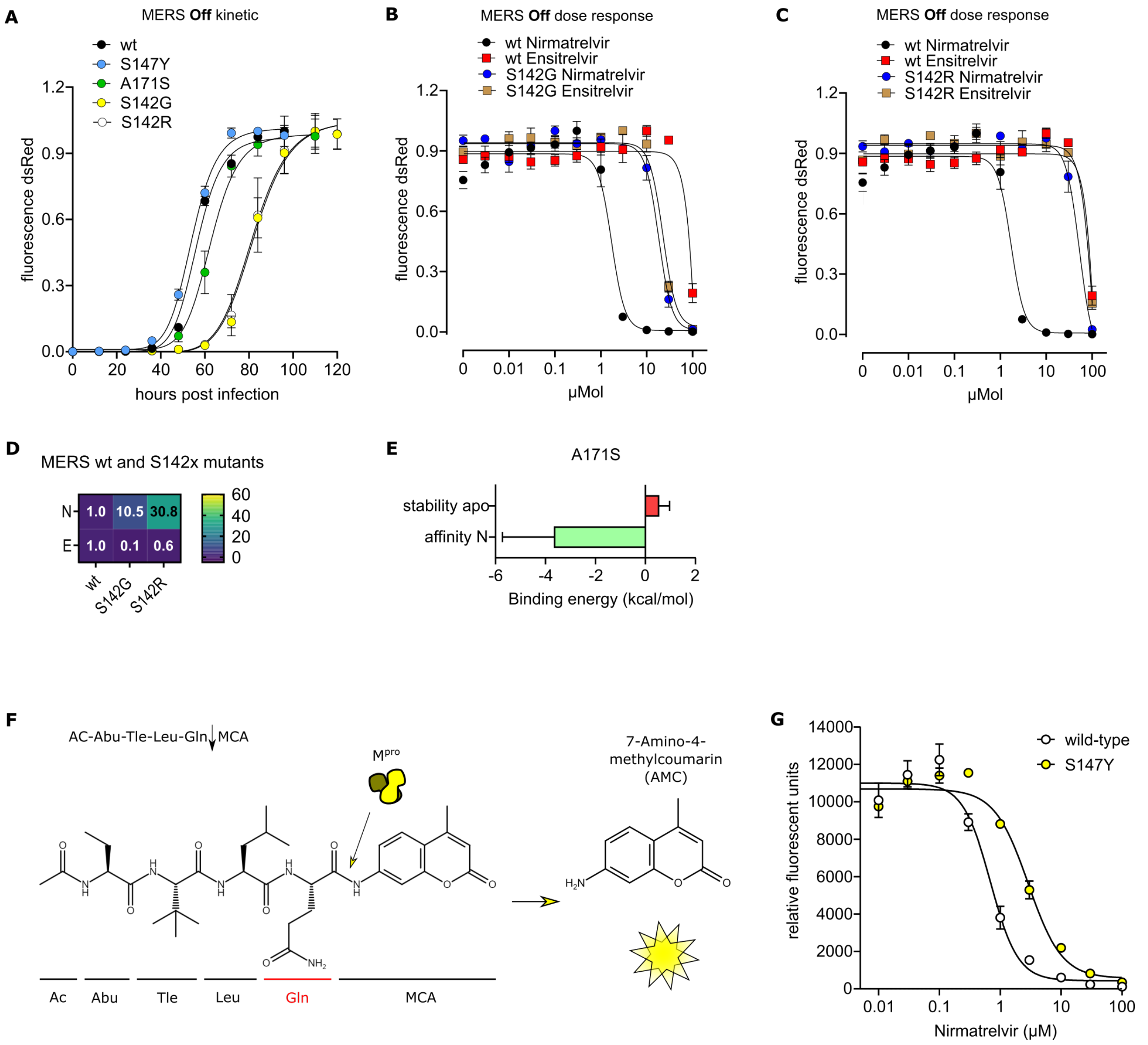

**Figure S6. Characterization of nirmatrelvir selected MERS-CoV-M$^{pro}$ mutants.** (**A**) M$^{pro}$-Off replication kinetics of MERS-CoV-M$^{pro}$ wt and nirmatrelvir selected MERS-CoV-M$^{pro}$ mutants S147Y, A171S, S142G and S142R. Data are presented as means of n = 8 biological replicates per condition. M$^{pro}$-Off assays to determine the susceptibility to nirmatrelvir (N) and ensitrelvir (E) in MERS-CoV-M$^{pro}$ wt and mutants S142G (**B**) and S142R (**C**). Data are presented as means of n = 4 biological replicates per condition. (**D**) Heat map showing $IC_{50}$ fold changes of (**B**) and (**C**). (**E**) Bar chart showing binding energies of A171S, indicating higher nirmatrelvir affinity and mildly reduced apo structure stability. (**F**) Fluorogenic substrate Ac-Abu-Tle-Leu-Gln-MCA cleavage by M$^{pro}$ releases 7-Amino-4-methylcoumarin (AMC). (**G**) Dose response of purified MERS-CoV-M$^{pro}$ proteins wt vs. S147Y mutant.

| | | 7alh | 7vh8 | 8dz0 | 7zqw | 5c3n | 7vtc | 3tlo | 5gwy | 6jij | 4dcd |
|---|---|---|---|---|---|---|---|---|---|---|---|
| **7alh** | apo-SARS2 | 0.00 | 0.53 | 0.72 | 0.48 | 0.88 | 0.74 | 0.90 | 0.89 | 0.79 | 2.26 |
| **7vh8** | NIR-SARS2 | 0.53 | 0.00 | 0.60 | 0.39 | 0.80 | 0.68 | 0.97 | 0.93 | 0.62 | 2.38 |
| **8dz0** | ENS-SARS2 | 0.72 | 0.60 | 0.00 | 0.65 | 0.75 | 0.72 | 0.95 | 1.01 | 0.81 | 2.40 |
| **7zqw** | AG7-SARS1 | 0.48 | 0.39 | 0.65 | 0.00 | 0.86 | 0.75 | 0.93 | 0.83 | 0.74 | 2.27 |
| **5c3n** | apo-MERS | 0.88 | 0.80 | 0.75 | 0.86 | 0.00 | 0.67 | 0.96 | 1.03 | 0.84 | 2.47 |
| **7vtc** | NIR_MERS | 0.74 | 0.74 | 0.72 | 0.75 | 0.67 | 0.00 | 0.93 | 0.87 | 0.76 | 2.24 |
| **3tlo** | apo-NL63 | 0.90 | 0.97 | 0.95 | 0.93 | 0.96 | 0.93 | 0.00 | 0.55 | 1.02 | 2.27 |
| **5gwy** | LEU-NL63 | 0.89 | 0.93 | 1.01 | 0.83 | 1.03 | 0.87 | 0.55 | 0.00 | 0.87 | 2.34 |
| **6jij** | LEU-MHV | 0.79 | 0.62 | 0.81 | 0.74 | 0.84 | 0.76 | 1.02 | 0.87 | 0.00 | 2.24 |
| **4dcd** | DIP-POLIO | 2.26 | 2.38 | 2.40 | 2.27 | 2.47 | 2.24 | 2.27 | 2.34 | 2.24 | 0.00 |

**Table S1.** Superposition matrix using residues 9 - 194 of chains A (inhibitor binding domain) for structures from different viruses. Secondary Structure Matching (SSM) superposition on Cα backbone atoms, RMSD in Å. As expected, global atomic RMSD differences between apo and inhibitor-bound forms of the same virus are generally smaller than differences between structures of distant viruses. The differences in inhibitor binding require a detailed local analysis of binding pocket conformation and ligand poses based on experimental structures or virtual docking results.

| sample | Position nct | Position aa | Codon wt | Codon mut | substitution | % snp | Localization |
|---|---|---|---|---|---|---|---|
| 1 | 865 | 289 | GAT | TAT | D289Y | 6.0% | allosteric |
| 1 | 403 | 135 | CCG | ACG | P135T | 86.5% | allosteric |
| 1 | 887 | 296 | GTA | GGA | V296G | 68.3% | allosteric |
| 2 | 616 | 206 | AAT | GAT | N206D | 95.3% | allosteric |
| 3 | 617 | 206 | AAT | ACT | N206T | 94.9% | allosteric |
| 4 | 875 | 292 | ACG | AAG | T292K | 18.7% | allosteric |
| 4 | 622 | 208 | GTC | TTC | V208F | 64.9% | allosteric |
| 5 | 865 | 289 | GAT | TAT | D289Y | 88.2% | allosteric |
| 6 | 797 | 266 | CTT | CCT | L266P | 92.6% | allosteric |
| 8 | 616 | 206 | AAT | GAT | N206D | 98.2% | allosteric |
| 9 | 625 | 209 | GCC | ACC | A209T | 78.4% | allosteric |
| 9 | 700 | 234 | AAT | TAT | N234Y | 9.1% | allosteric |
| 9 | 362 | 121 | TAC | TTC | Y121F | 60.2% | near catalytic site |
| 10 | 698 | 233 | TTC | TCC | F233S | 93.9% | allosteric |
| 11 | 618 | 206 | AAT | AAG | N206K | 88.6% | allosteric |
| 12 | 734 | 245 | TTT | TCT | F245S | 9.7% | allosteric |
| 12 | 617 | 206 | AAT | ACT | N206T | 45.7% | allosteric |
| 12 | 878 | 293 | CCG | CAG | P293Q | 7.6% | allosteric |
| 12 | 424 | 142 | AGC | GGC | S142G | 89.0% | catalytic site |
| 13 | 697 | 233 | TTC | GTC | F233V | 4.6% | allosteric |
| 13 | 618 | 206 | AAT | AAA | N206K | 14.2% | allosteric |
| 13 | 634 | 212 | TAC | CAC | Y212H | 19.5% | allosteric |
| 13 | 899 | 300 | ATA | AAA | I300K | 97.7% | cleavage site |
| 13 | 8 | 3 | TTG | TCG | L3S | 48.4% | cleavage site |
| 14 | 699 | 233 | TTC | TTA | F233L | 91.6% | allosteric |
| 15 | 626 | 209 | GCC | GAC | A209D | 8.0% | allosteric |
| 15 | 641 | 214 | GCG | GAG | A214E | 50.7% | allosteric |
| 15 | 616 | 206 | AAT | CAT | N206H | 8.1% | allosteric |
| 15 | 618 | 206 | AAT | AAG | N206K | 20.0% | allosteric |
| 15 | 424 | 142 | AGC | CGC | S142R | 33.0% | catalytic site |
| 16 | 887 | 296 | GTA | GGA | V296G | 87.8% | allosteric |
| 17 | 775 | 259 | GGG | TGG | G259W | 10.6% | allosteric |
| 17 | 778 | 260 | GTT | TTT | V260F | 96.4% | allosteric |
| 17 | 706 | 236 | TGG | GGG | W236G | 38.1% | allosteric |
| 17 | 556 | 186 | GGT | TGT | G186C | 7.0% | near catalytic site |
| 18 | 865 | 289 | GAT | AAT | D289N | 7.2% | allosteric |
| 18 | 787 | 263 | GAG | AAG | E263K | 3.7% | allosteric |
| 18 | 718 | 240 | AAC | GAC | N240D | 97.5% | allosteric |
| 18 | 683 | 228 | ACT | AAT | T228N | 75.8% | allosteric |
| 19 | 607 | 203 | TGT | CGT | C203R | 84.4% | allosteric |
| 19 | 706 | 236 | TGG | CGG | W236R | 6.6% | allosteric |
| 20 | 632 | 211 | CTC | CCC | L211P | 7.7% | allosteric |
| 20 | 896 | 299 | CAG | CCG | Q299P | 57.9% | allosteric |
| 21 | 787 | 263 | GAG | AAG | E263K | 16.4% | allosteric |
| 21 | 779 | 260 | GTT | GGT | V260G | 62.9% | allosteric |

| | | | | | | | |
|---|---|---|---|---|---|---|---|
| 21 | 71 | 24 | AGC | ATC | S24I | 8.1% | near catalytic site |
| 22 | 697 | 233 | TTC | CTC | F233L | 44.8% | allosteric |
| 22 | 598 | 200 | GAC | TAC | D200Y | 28.9% | near catalytic site |
| 23 | 464 | 155 | AAG | ACG | K155T | 11.1% | allosteric |
| 23 | 616 | 206 | AAT | GAT | N206D | 22.5% | allosteric |
| 23 | 886 | 296 | GTA | TTA | V296L | 95.5% | allosteric |
| 23 | 17 | 6 | ATG | AAG | M6K | 6.0% | cleavage site |
| 23 | 556 | 186 | GGT | AGT | G186S | 8.1% | near catalytic site |
| 24 | 625 | 209 | GCC | ACC | A209T | 32.5% | allosteric |
| 24 | 734 | 245 | TTT | TCT | F245S | 15.7% | allosteric |
| 24 | 617 | 206 | AAT | AGT | N206S | 7.6% | allosteric |
| 24 | 610 | 204 | TCT | CCT | S204P | 37.6% | allosteric |
| 24 | 779 | 260 | GTT | GGT | V260G | 10.6% | allosteric |
| 24 | 241 | 81 | GTG | ATG | V81M | 5.4% | allosteric |
| 24 | 899 | 300 | ATA | AAA | I300K | 97.7% | cleavage site |
| 24 | 599 | 200 | GAC | GGC | D200G | 13.0% | near catalytic site |
| 25 | 625 | 209 | GCC | ACC | A209T | 6.7% | allosteric |
| 25 | 865 | 289 | GAT | AAT | D289N | 20.4% | allosteric |
| 25 | 628 | 210 | TGG | CGG | W210R | 92.2% | allosteric |
| 26 | 866 | 289 | GAT | GGT | D289G | 17.7% | allosteric |
| 26 | 787 | 263 | GAG | AAG | E263K | 5.8% | allosteric |
| 26 | 617 | 206 | AAT | ACT | N206T | 22.3% | allosteric |
| 26 | 668 | 223 | GTA | GGA | V223G | 12.4% | allosteric |
| 26 | 887 | 296 | GTA | GCA | V296A | 10.0% | allosteric |
| 26 | 424 | 142 | AGC | GGC | S142G | 78.0% | catalytic site |
| 27 | 625 | 209 | GCC | ACC | A209T | 24.4% | allosteric |
| 27 | 698 | 233 | TTC | TGC | F233C | 57.0% | allosteric |
| 27 | 616 | 206 | AAT | GAT | N206D | 23.7% | allosteric |
| 28 | 788 | 263 | GAG | GGG | E263G | 7.1% | allosteric |
| 28 | 775 | 259 | GGG | TGG | G259W | 94.0% | allosteric |
| 29 | 880 | 294 | GAG | AAG | E294K | 7.5% | allosteric |
| 29 | 618 | 206 | AAT | AAG | N206K | 80.4% | allosteric |
| 30 | 616 | 206 | AAT | GAT | N206D | 98.4% | allosteric |
| 31 | 616 | 206 | AAT | GAT | N206D | 86.1% | allosteric |
| 31 | 424 | 142 | AGC | GGC | S142G | 97.0% | catalytic site |
| 32 | 797 | 266 | CTT | CCT | L266P | 64.5% | allosteric |
| 32 | 887 | 296 | GTA | GGT | V296G | 6.5% | allosteric |
| 33 | 617 | 206 | AAT | ACT | N206T | 77.2% | allosteric |
| 33 | 772 | 258 | ACA | CCA | T258P | 9.3% | allosteric |
| 34 | 625 | 209 | GCC | ACC | A209T | 11.4% | allosteric |
| 34 | 709 | 237 | GCC | ACC | A237T | 8.1% | allosteric |
| 34 | 803 | 268 | GCC | CCC | A268P | 23.2% | allosteric |
| 34 | 617 | 206 | AAT | AGT | N206S | 14.2% | allosteric |
| 34 | 611 | 204 | TCT | TTT | S204F | 29.2% | allosteric |
| 35 | 734 | 245 | TTT | TCT | F245S | 95.7% | allosteric |
| 36 | 625 | 209 | GCC | ACC | A209T | 10.0% | allosteric |

| | | | | | | | |
|---|---|---|---|---|---|---|---|
| 36 | 866 | 289 | GAT | GGT | D289G | 71.9% | allosteric |
| 36 | 787 | 263 | GAG | AAG | E263K | 14.3% | allosteric |
| 36 | 418 | 140 | AAG | GAG | K140E | 9.5% | near catalytic site |
| 37 | 697 | 233 | TTC | GTC | F233V | 15.0% | allosteric |
| 37 | 734 | 245 | TTT | TCT | F245S | 14.9% | allosteric |
| 37 | 480 | 160 | ATT | ATG | I160M | 15.3% | allosteric |
| 37 | 616 | 206 | AAT | GAT | N206D | 24.6% | allosteric |
| 37 | 618 | 206 | AAT | AAG | N206K | 9.1% | allosteric |
| 37 | 886 | 296 | GTA | TTA | V296L | 11.0% | allosteric |
| 37 | 418 | 140 | AAG | GAG | K140E | 7.1% | near catalytic site |
| 38 | 401 | 134 | CGA | CAA | R134Q | 96.9% | allosteric |
| 39 | 806 | 269 | ATA | AGA | I269R | 49.1% | allosteric |
| 39 | 887 | 296 | GTA | GGA | V296G | 39.1% | allosteric |
| 40 | 866 | 289 | GAT | GGT | D289G | 9.7% | allosteric |
| 40 | 698 | 233 | TTC | TCC | F233S | 79.8% | allosteric |
| 41 | 637 | 213 | GCC | ACC | A213T | 27.6% | allosteric |
| 41 | 863 | 288 | GAA | GGA | E288G | 90.3% | allosteric |
| 41 | 335 | 112 | GGC | GAC | G112D | 10.2% | allosteric |
| 41 | 31 | 11 | GGT | AGT | G11S | 22.3% | allosteric |
| 42 | 887 | 296 | GTA | GGA | V296G | 85.1% | allosteric |
| 43 | 616 | 206 | AAT | GAT | N206D | 78.7% | allosteric |
| 44 | 634 | 212 | TAC | GAC | Y212D | 96.3% | allosteric |
| 45 | 866 | 289 | GAT | GGT | D289G | 90.3% | allosteric |
| 45 | 622 | 208 | GTC | TTC | V208L | 5.9% | allosteric |
| 45 | 511 | 171 | GCG | TCG | A171S | 31.6% | catalytic site |
| 46 | 865 | 289 | GAT | TAT | D289Y | 83.2% | allosteric |
| 46 | 616 | 206 | AAT | GAT | N206D | 6.6% | allosteric |
| 46 | 718 | 240 | AAC | GAC | N240D | 97.0% | allosteric |
| 47 | 787 | 263 | GAG | AAG | E263K | 76.4% | allosteric |
| 47 | 887 | 296 | GTA | GCA | V296A | 16.5% | allosteric |
| 47 | 557 | 186 | GGT | GTT | G186V | 7.8% | near catalytic site |
| 47 | 634 | 121 | TAC | AAC | Y121N | 5.9% | near catalytic site |
| 48 | 700 | 234 | AAT | TAT | N234Y | 93.2% | allosteric |
| 48 | 564 | 188 | TTT | TTG | F188L | 27.2% | catalytic site |
| 48 | 440 | 147 | TCC | TAC | S147Y | 13.4% | catalytic site |

**Table S2. Mutations selected with recombinant VSV-MERS-$M^{pro}$.** Columns: position nucleotide (nct), position amino acid (aa), codon wild-type (wt), codon mutant (mut), amino acid substitution, percentage of the mutation / single nucleotide polymorphism within each well (% snp), localization of the substitution.

| Name | Sequence (5'-3' direction) |
|---|---|
| **VSV-G-M$^{pro}$-L** | |
| G-33n-before-KpnI-for | GAACCGGTCCTGCTTTCACC |
| G-rev | CTTTCCAAGTCGGTTCATCTC |
| G-cut1-7aa-for | GAGATGAACCGACTTGGAAAGATCACTAGCGGTGTATTGCAG |
| cut2-7aa-L-rev | GTCTCAAAATCGTGGACTTCCATTGTTACTTTTCTTACACCGGACTG |
| G-cut1-8aa-for | GAGATGAACCGACTTGGAAAGTCAATCACTAGCGGTGTATTGCAG |
| cut2-8aa-L-rev | GTCTCAAAATCGTGGACTTCCATGTATGTTACTTTTCTTACACCGGACTG |
| L-for | ATGGAAGTCCACGATTTTGAGACCGACG |
| L-33n-after-HpaI-rev | ATGGAAGTCCACGATTTTGAGACCGACG |
| **MERS-M$^{pro}$-On-Nt-QtoN** | |
| Hygro-P-for | CTGTTTTGACCTCCATAGAAGATTCTAGAGCTAGCATGGATAATCTCACAAAAGTTC |
| P-GGSG-rev | GCTCCCTCCGCCGCTTCCGCCATCTGATACTGCTTCTGATTGG |
| MERS-On-N-term-QtoN-for | GGCGGAAGCGGCGGAGGGAGCGGGGGCGGGAGCGGATCAATCACTAGCGGTGTATTGAACAGTGGTTTGGTC |
| MERS-On-C-term-rev | GCCGGATCCACCGCCTGAGCCGCCTCCGGACCCTCCGTATGTTACTTTTCTTACACCGGAC |
| GGSG-P-for | GGCTCAGGCGGTGGATCCGGCGTTTGGTCTCTCTCAAAGACAT |
| Hygro-P-rev | GAGGGAGAGGGGCGGATCCCCTTAATTAACTACAGAGAATATTTGACTCTCGC |
| **MERS-M$^{pro}$-Off** | |
| Blasti-L-for | CTGTTTTGACCTCCATAGAAGATTCTAGAGCTAGCATGGAAGTCCACGATTTTGAG |
| L-blasti-rev | GAGGGAGAGGGGCGGATCCCCTTAATTAATTAATCTCTCCAAGAGTTTTCCTC |
| Blasti-for | CATTCGATTAGTGAACGGATCTC |
| GFP-rev | CTTGTACAGCTCGTCCATGCC |
| GFP-cut1-MERS-Off-7aa-for | GGCATGGACGAGCTGTACAAGATCACTAGCGGTGTATTGCAG |
| 7aa-MERS-Off-cut2-L-rev | GTCTCAAAATCGTGGACTTCCATTGTTACTTTTCTTACACCGGACTG |
| L-for | ATGGAAGTCCACGATTTTGAGACCGACG |
| L-33n-after-HpaI-rev | GATGTTGGGATGGGATTGGC |
| **Mutation primers** | |
| SARS-2-T25A-for | CAAGTAACTTGTGGTACAGCTACAC |
| SARS-2-T25A-rev | GACCGTTAAGTGTAGCTGTACC |
| SARS-2-T25N-for | CAAGTAACTTGTGGTACAAATACACTTAACG |
| SARS-2-T25N-rev | GCCAAAGACCGTTAAGTGTATTTGTAC |
| SARS-2-T25M-for | CAAGTAACTTGTGGTACAATGACACTTAAC |
| SARS-2-T25M-rev | GACCGTTAAGTGTCATTGTACCAC |
| MERS-M25T-for | CCTGTGGTAGCACTACACTTAATG |
| MERS-M25T-rev | CCAGAGTCCATTAAGTGTAGTGCTAC |

| | |
|---|---|
| MERS-S142G-for | CGAATTATACGATTAAGGGTGGCTTTTTG |
| MERS-S142G-rev | CCACACAAAAAGCCACCC |
| MERS-S142R-for | CGAATTATACGATTAAGGGTCGCTTTTTG |
| MERS-S142R-rev | CCACACAAAAAGCGACCC |
| MERS-S147Y-for | GTAGCTTTTTGTGTGGTTACTGTGG |
| MERS-S147Y-rev | CGACAGAACCACAGTAACCAC |
| MERS-A171S-for | CACCAAATGGAACTCTCGAACG |
| MERS-A171S-rev | GGTATGAGTACCGTTCGAGAGTTC |
| MERS-S24T-for | CAAGTAACCTGTGGTACAATGACAC |
| MERS-S24T-rev | GAGTCCATTAAGTGTCATTGTACCAC |
| MERS-S24T-M25T-for | CCAAGTAACCTGTGGTACAACTACAC |
| MERS-S24T-M25T-rev | CCAGAGTCCATTAAGTGTAGTTGTACC |
| **Other On constructs** | |
| SARS-2-On-N-term-QtoN-for | CACCTCAGCTGTTTTGAACAGTGG |
| SARS-2-On-N-term-QtoN-rev | CTAAAACCACTGTTCAAAACAGC |
| SARS-1-On-N-term-QtoN-for | GGCGGAAGCGGCGGAGGGAGCGGGGGCGGAGCGGAAGTA TCACGTCTGCTGTGCTCAACTCAGGCTTCAG |
| SARS-1-GGSG-On-rev | GCCGGATCCACCGCCTGAGCCGCCTCCGGACCCTCCTTTGAC TATTTTTTTGAACTTACCTTG |
| HKU9-On-N-term-QtoN-for | GGCGGAAGCGGCGGAGGGAGCGGGGGCGGAGCGGAAGCG TCGCCAGTGCTGCGCTCAACGCGGGTCTTACTC |
| HKU9-GGSG-On-rev | GCCGGATCCACCGCCTGAGCCGCCTCCGGACCCTCCTCGAAA CATAGATTGAAATTTACCTTG |
| HCoV-NL63-On-N-term-QtoN-for | GGCGGAAGCGGCGGAGGGAGCGGGGGCGGAGCGGAATCA GTTACAATAGTACCTTGAACAGCGGACTG |
| HCoV-NL63-GGSG-On-rev | GCCGGATCCACCGCCTGAGCCGCCTCCGGACCCTCCAAGCCC GAATATAACCTTTCCTG |
| HCoV-229E-On-N-term-QtoN-for | GGCGGAAGCGGCGGAGGGAGCGGGGGCGGAGCGGAGTAT CTTATGGCTCAACGCTCAACGCCGGCTTGCGC |
| HCoV-229E-GGSG-On-rev | GCCGGATCCACCGCCTGAGCCGCCTCCGGACCCTCCAAACAT GGATGTAGTCTTACCAGATTG |
| MHV-A59-On-N-term-QtoN-for | GGCGGAAGCGGCGGAGGGAGCGGGGGCGGAGCGGATCAG TCACCACTTCATTTCTCAACTCCGGGATAG |
| MHV-A59-GGSG-On-rev | GCCGGATCCACCGCCTGAGCCGCCTCCGGACCCTCCTTTTAT TACTCTAGTCCTTTTACTCTGCAG |
| Polio-On-N-term-QtoN-for | GGCGGAAGCGGCGGAGGGAGCGGGGGCGGAGCGGAACCA TTCGGACAGCAAAGGTAAACGGACCAGGGTTC |
| Polio-GGSG-On-rev | GCCGGATCCACCGCCTGAGCCGCCTCCGGACCCTCCAGGTCT CATCCACTGGATTTC |
| ***E.coli*** **$M^{pro}$ expression plasmid** | |
| $M^{pro}$-expr.-S147Y-for | GCTTTCTGTGTGGTTACTGCGG |
| $M^{pro}$-expr.-S147Y-rev | CGCTACCGCAGTAACCACAC |

**Table S3.** Cloning oligonucleotides / primers.

**Alignments S1 to S4**

Nucleotide sequence alignments of SARS-CoV-2 and -1 main proteases; SARS-CoV-2 and MERS-CoV main proteases; SARS-CoV-2 and -1 spike; SARS-CoV-2 and MERS-CoV spike in fasta alignment format.

**Plasmids S1 to S7**

Plasmid S1: VSV-Indiana antigenome pBluescript expression plasmid under a T7 polymerase promoter and terminator with hepatitis-D ribozyme and ampicillin resistance.

Plasmid S2: VSV-Indiana plasmid with insertion of the MERS-CoV $M^{pro}$ and cognate cleavage sites replacing the VSV intergenic region between VSV-G and VSV-L.

Plasmid S3: Lentiviral expression vector modified from Addgene pLenti CMVie-IRES-BlastR accession #119863 by replacing blasticidin resistance with hygromycin resistance to generate pLenti CMVie-IRES-HygroR.

Plasmid S4: pLenti CMVie-IRES-HygroR encoding VSV-P with intramolecular insertion of MERS-CoV $M^{pro}$. N-terminal glutamine was mutated to asparagine to knock-out one of the two cis-cleavage sites of $M^{pro}$. VSV-P with intramolecular insertion of MERS-CoV $M^{pro}$ constitutes $M^{pro}$-On construct.

Plasmid S5: pLenti CMVie-IRES-BlastR encoding VSV-L polymerase gene.

Plasmid S6: pLenti CMVie-IRES-BlastR encoding VSV-L polymerase gene with a N-terminal tag of MERS-CoV $M^{pro}$ and GFP, resulting in $M^{pro}$-Off construct.

Plasmid S7: E.coli MERS-CoV-Mpro expression plasmid with N-terminal His-tag and cleavable sequence for His-tag removal.

**Sequences S1 to S8**

Sequence S1-8: SARS-CoV-2, SARS-CoV-1, MERS-CoV, HKU9, NL63, 229E and MHV main proteases and poliovirus 3C-protease sequences as they were used in On, Off and virus constructs.
